# Distinct cortical patches for syntactic and semantic composition in the human brain

**DOI:** 10.64898/2026.06.18.732834

**Authors:** Thomas Dighiero-Brecht, Naama Friedmann, Luigi Rizzi, Christophe Pallier, Stanislas Dehaene

## Abstract

Although the brain areas for language processing are well delimited, whether lexical-semantic and syntactic processes are spatially segregated remains debated. To clarify this issue, we conducted two experiments using 7-Tesla functional MRI in 20 participants performing: a functional localizer involving reading sequences of words of increasing linguistic complexity; and a presentation of short, semantically impoverished three-word mini-sentences, flashed in a single glance (e.g., “he does it”), whose grammaticality and syntactic complexity was manipulated through syntactic movement. Our results reveal two functionally dissociable sets of cortical patches within the language system: one sensitive to syntactic structure even in the absence of meaning, and the other involved in semantic composition. This dual-network architecture was consistently observed in the majority of participants, although its precise anatomical localization varied. The two types of voxels coexisted even within a given brain region of the Glasser atlas. Results were confirmed using subject-specific analyses and region-by-condition interactions, as voxels in those two systems displayed markedly different responses to mini-sentences. Thus, high-resolution functional imaging reveals a division of labor between syntactic and semantic composition within the classical language network.

## Introduction

The human capacity for language is a defining feature of our species, allowing us to convey complex thoughts and abstract ideas with remarkable efficiency. This virtually unlimited expressive power stems from the compositional nature of language, where meaning is conveyed not only by individual lexical items, but also by the specific rules governing their arrangement (Chomsky, 1965). At the brain level, language processing involves a robust network of fronto-temporal areas, typically lateralized to the left hemisphere, known as the "language network". This system is characterized by high intra- and inter-individual reproducibility and a striking functional specificity for linguistic inputs over other complex cognitive domains, such as music or mathematics (Fedorenko et al., 2024). However, the exact computational architecture underlying this network remains a subject of intense investigation. A classic hypothesis proposes a functional division of labor, with a subset of regions dedicated to processing formal grammatical structure (*syntax*) and others specialized for the computation of meaning (*semantics*). Evidence for syntactic and semantic hubs has been reported across various studies, though their precise anatomical boundaries varied (Bornkessel-Schlesewsky & Schlesewsky, 2013; Duffau et al., 2014; Frankland & Greene, 2015; Hagoort, 2005; Hickok & Poeppel, 2007; Ralph et al., 2017; Tyler et al., 2008, 2010, 2011). Syntactic hubs were theoretically defined by their sensitivity to the abstract, hierarchical constituent structures of language, often modeled as a linguistic tree (Dehaene et al., 2015; Pattamadilok et al., 2016), and their activation independently of whether the resulting syntactic construction is meaningful or not (Goucha & Friederici, 2015; Mazoyer et al., 1993; Pallier et al., 2011; Woolnough et al., 2023). Some of the evidence for this segregation builds upon the work of Pallier et al. (2011), who demonstrated that some posterior temporal and inferior frontal regions showed an activation proportional to the complexity of constituent structures and which was equivalent for both real-word and "Jabberwocky" sentences (whose content words were replaced with pseudowords, rendering them meaningless). This finding suggested that a core set of brain areas, particularly in posterior superior temporal sulcus and inferior frontal gyrus, is involved in syntactic constituent building, independently of lexical meaning.

However, the hypothesis of a cortical specialization has been challenged by researchers advocating for a more distributed sensitivity, where syntax and lexico-semantic processes would be integrated across much of the language network. Recently, Shain et al. (2024) sought to replicate the Pallier et al. (2011) findings using contemporary methods of 3-Tesla functional MRI, including a full within-subject design and functional localizers permitting the use of independent data for within-subject ROI definition. While they successfully replicated the overall effect of increased activation in the language areas as a function of constituent size, as well as the finding that this pattern continues to hold in several areas even when lexical content is removed (in Jabberwocky), they claimed that no region met the criteria for a purely syntactic hub, since all showed an effect of lexicality, with stronger responses to real-word sentences than to Jabberwocky. These results were taken as congruent with the hypothesis of a lack of syntax-semantics segregation within the macro-anatomical language network (Fedorenko et al., 2020, 2024).

Note however that, in theory, the finding of a lexicality effect (greater response to stimuli made of real words than of pseudowords) need not imply that a region is solely involved in semantics. This is because different components of the mental lexicon convey not only semantic but also syntactic features (such as part-of-speech, grammatical gender, number of arguments for verbs and their thematic roles, and more, Biran & Friedmann, 2012; Katz & Friedmann, 2024). Unless great care is taken to preserve them, many if not all of these features vanish or become opaque in Jabberwocky sentences, jeopardizing syntactic computations and potentially causing a reduced activation in both syntactic and semantic networks. This logic may explain Shain et al.’s (2024) additional observation of a lexical by constituent size interaction in several areas: as constituent size increases, the missing syntactic cues in Jabberwocky may increasingly impede the construction of a full syntactic tree. It is with this critique in mind that, in the present work, we did not use Jabberwocky, but studied the brain response to syntactically correct yet semantically anomalous sentences, analogous to Chomsky’s famous “colorless green ideas sleep furiously”. We reasoned that cortical patches involved in syntax should respond identically to such stimuli and to normal meaningful sentences, and more so than to lists of words.

Beyond the challenges linked to Jabberwocky, other neurolinguistic evidence continues to support the existence of specialized structural processing. In modern linguistic theory, following pioneering work from Chomsky and colleagues, the "Merge" operation is postulated to serve as the fundamental mechanism for sentence construction (Rizzi, 2012). Such merge can be separated into external merge, i.e., stitching together phrases to form more complex and semantically richer phrases or sentences, and internal merge, i.e., changing the position of words or groups of words in a given sentence to create operator-variable structures and other configurations which impact on the intended meaning. Thus, internal merge is often referred to as syntactic movement, which can take several forms depending on the characteristics and nature of the displaced structure. Previous studies have looked for and identified neural correlates of such syntactic operations. In a pioneer study, Ben-Shachar and colleagues identified left-hemispheric posterior temporal and inferior frontal regions involved in syntactic movement, namely topicalization, and wh-questions in Hebrew (Ben-Shachar et al., 2004). Another study by Shetreet and Friedmann found analogous results regarding wh movement using a different set of Hebrew sentences and participants (Shetreet & Friedmann, 2014). Shetreet and Friedmann also reported separate networks dedicated to wh movement and verb movement. More recently, Park et al. also identified brain regions involved in wh-constructions in Korean, which share the same general involvement of posterior temporal and inferior frontal regions of the left hemisphere (Park et al., 2024). Further evidence from pre- and post-surgical mapping of fine-grained language impairment reinforces the claim that some regions are indeed critical for syntax (Kahana et al., 2025).

Here, we argue that one of the main obstacles in resolving this debate is the spatial resolution limit of 3 Tesla fMRI. Recent intracranial recordings demonstrate that electrodes separated by only a few millimeters can assume distinct language-related computations (Fedorenko et al., 2016; Murphy et al., 2022; Nelson et al., 2017; Regev et al., 2024; Woolnough et al., 2020, 2023). This suggests that specialization exists at a scale beneath that of standard non-invasive neuroimaging techniques, and may require access to a mesoscopic scale cortical organization (∼1 mm and below). Single-subject ultra-high-field fMRI, at 7 tesla and above, has already begun to transform our understanding of cortical specialization (Allen et al., 2021; Cai et al., 2023). For instance, the Visual Word Form Area (VWFA), long viewed as a single spatially contiguous region, has been revealed by 7T imaging to be a fine-grained "archipelago" of millimetric cortical patches with varying profiles of response to different visual categories and languages in bilinguals (Zhan et al., 2023). Applying this individual, high-resolution yet whole-brain approach to the entire language network offers the potential to bridge the gap between macroscopic network views and the microscopic reality of neuronal specialization.

In the present study, we designed a series of two 7T fMRI experiments in the same volunteers to ask whether high resolution single-subject analysis allows us to detect fine-grained distinctions within the language network. Experiment 1 was designed as a classical sentence-based subject-specific language localizer, in accordance with state-of-the-art methodology in language fMRI studies (Fedorenko et al., 2010; Nieto-Castañón & Fedorenko, 2012), with ∼9-word stimuli presented in a Rapid Serial Visual Presentation (RSVP) and including contrasts capable of separating putative syntactic and semantic voxels (see below). Experiment 2 then sought to confirm whether these voxels showed distinct response to syntactic movement in semantically impoverished mini-sentences made of only 3 words flashed all at once on screen.

We designed the experiment 1 localizer by adapting the visual MathLang paradigm designed in our lab, which consists in rating the truth value of sequentially presented written statements (or non-sensical word sequences) including mathematical and non-mathematical sentences matched for low-level linguistic features (Amalric & Dehaene, 2016; Moreno et al., 2025). For present purposes, we excluded the math conditions and presented only a hierarchy of five types of word sequences ranging from lists of consonant strings to word lists, semantically anomalous sentences, and two types of meaningful sentences.

For experiment 2, we flashed extremely short sentences with limited semantic content, whose word-size length was short enough to be read in a single glance. We use a Rapid *Parallel* Visual Presentation (RPVP), which recently revealed a capacity for multiword integration within a single fixation in both magnetoencephalography and behavior (Flower & Pylkkänen, 2024; Snell & Grainger, 2017). The stimuli were in French and comprised a pronoun, a short verb and either a second pronoun or a locative (e.g. tu le vois [you see it], tu vas là [you go there]), in various orders. Those stimuli were highly impoverished in their semantic content, but syntactically rich, and we further manipulated syntactic complexity by adding various forms of movement (clitic, verb, and wh-movement).

To anticipate on the results, experiment 1 identified distinct sets of voxels involved in syntactic and semantic processing. At the individual level, these two functional populations were found to coexist within most language areas of the Glasser atlas, suggesting a much finer-grained internal organization than the macroscopic regions typically described. These results were validated through cross-validation and replication in an independent cohort. Experiment 2 further corroborated this dissociation by revealing that the two types of voxels exhibit very different profiles of activation to the semantically impoverished mini-sentences. Specifically, in syntax-related voxels, activation increased with syntactic movement, whereas a deactivation was found in semantic voxels.

## Results

### Distinct Cortical Networks for Syntactic and Semantic Processing

In experiment 1, to separate the putative neural substrates of syntax and semantics, we employed a hierarchical Rapid Serial Visual Presentation (RSVP) truth-judgment task (Fig. 1A). The results are presented in Figure 2. In the primary cohort (n=20), at the group level, semantically anomalous but syntactically well-formed sentences, compared to unstructured word lists, elicited robust, widespread activation within the "core language network" (p<.001, FDR-corrected at α<.05; Figure 2A). In these results, we refer to these as “syntax voxels” purely for simplicity and solely to refer to the contrast used to isolate them (we return to their interpretation further below). Significant clusters of syntax voxels were identified along the superior temporal gyrus (STG), middle frontal gyrus (MFG), and inferior frontal gyrus (IFG). At the individual level (Figure 2B), prevalence analysis demonstrated high anatomical consistency across participants: activation was observed in the anterior STG (aSTG) for 100% of subjects (20/20), the posterior STG (pSTG) in 80% (16/20), the MFG in 90% (18/20), and the IFG in 90% (18/20 for pars triangularis) to 100% (20/20 for pars orbitalis). These results were qualitatively and quantitatively replicated in an independent cohort (n=13), with comparable prevalence rates across all key regions (aSTG: 92%, pSTG: 100%, MFG: 92%, IFGtri: 85%, IFGorb: 85%) and a spatial distribution highly consistent with previous reports (Moreno et al., 2025).

**Figure 1:**
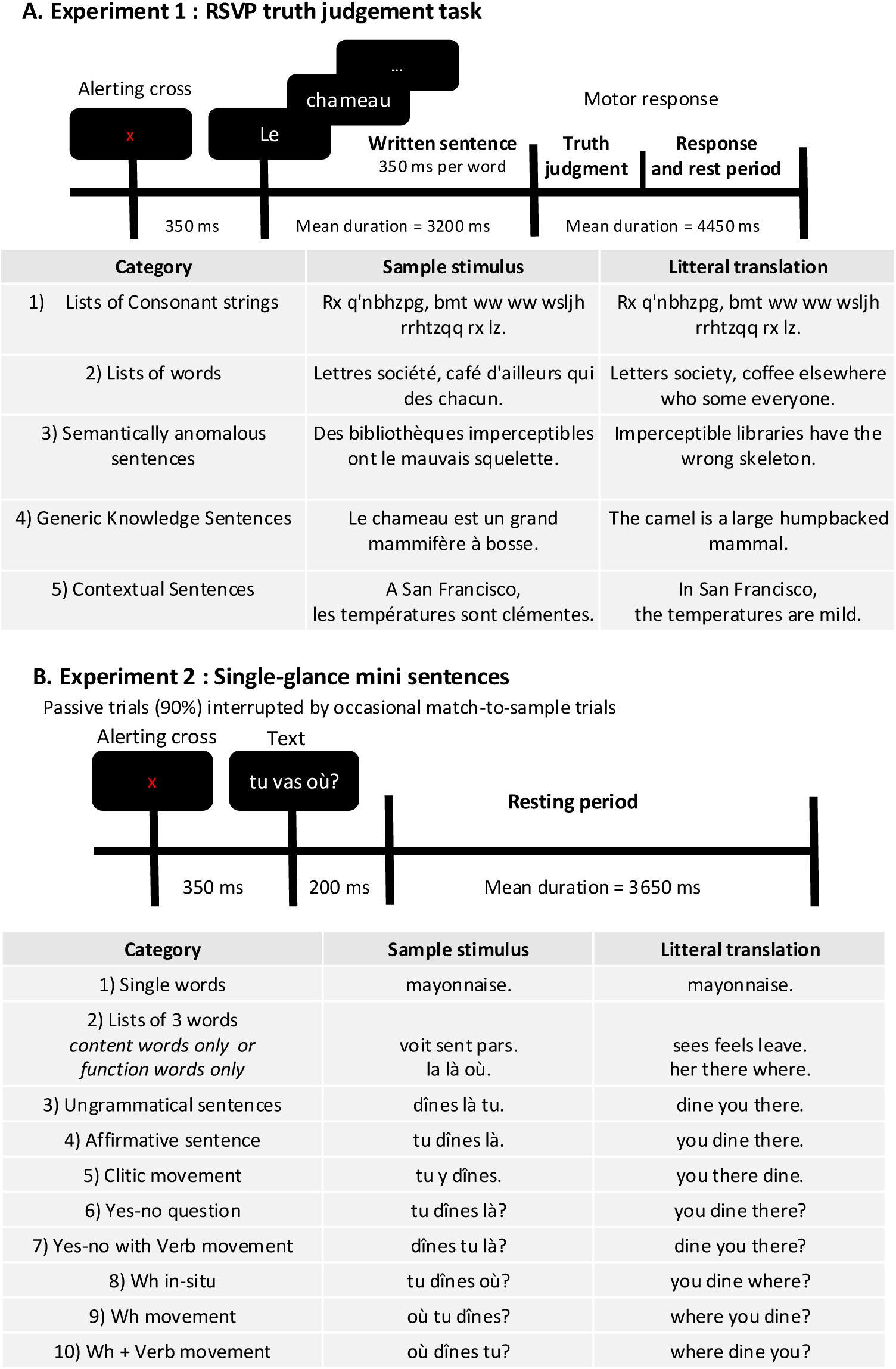
Experimental design and linguistic stimulus space. To isolate the neural correlates of syntactic and semantic processing, two complementary fMRI tasks were designed to probe the linguistic hierarchy at different temporal scales. A: RSVP truth judgement task (Task 1). Stimuli consisted of 100 sentences across five categories, presented using Rapid Serial Visual Presentation (RSVP; 350 ms per word). The design orthogonally manipulated three core linguistic factors: lexicality (Lex), syntax (Syn), and semantics (Sem). Conditions ranged from low-level controls (pseudo-word strings; Lex–, Syn–, Sem–) to fully formed factual sentences (Lex+, Syn+, Sem+). This hierarchy allows for the precise isolation of syntactic structure (Semantically anomalous sentences vs. Word lists) and semantic content (Generic knowledge sentences vs. Semantically anomalous sentences). At the end of each trial, participants performed a truth-value judgment to ensure high-level propositional processing. B: Single-glance mini-sentences (Task 2). To investigate the speed and automaticity of the language network, 240 stimuli across ten categories were presented as a single-glance display (200 ms duration). The stimulus set included three control conditions and seven grammatically correct sentence types with varying degrees of syntactic complexity. To maintain a focus on naturalistic, rapid processing, 90% of the trials were passive. Attentional engagement was monitored via occasional "catch trials" (10%), where participants performed a match-to-sample task, identifying which of two stimuli had been presented in the preceding trial.

**Figure 2:**
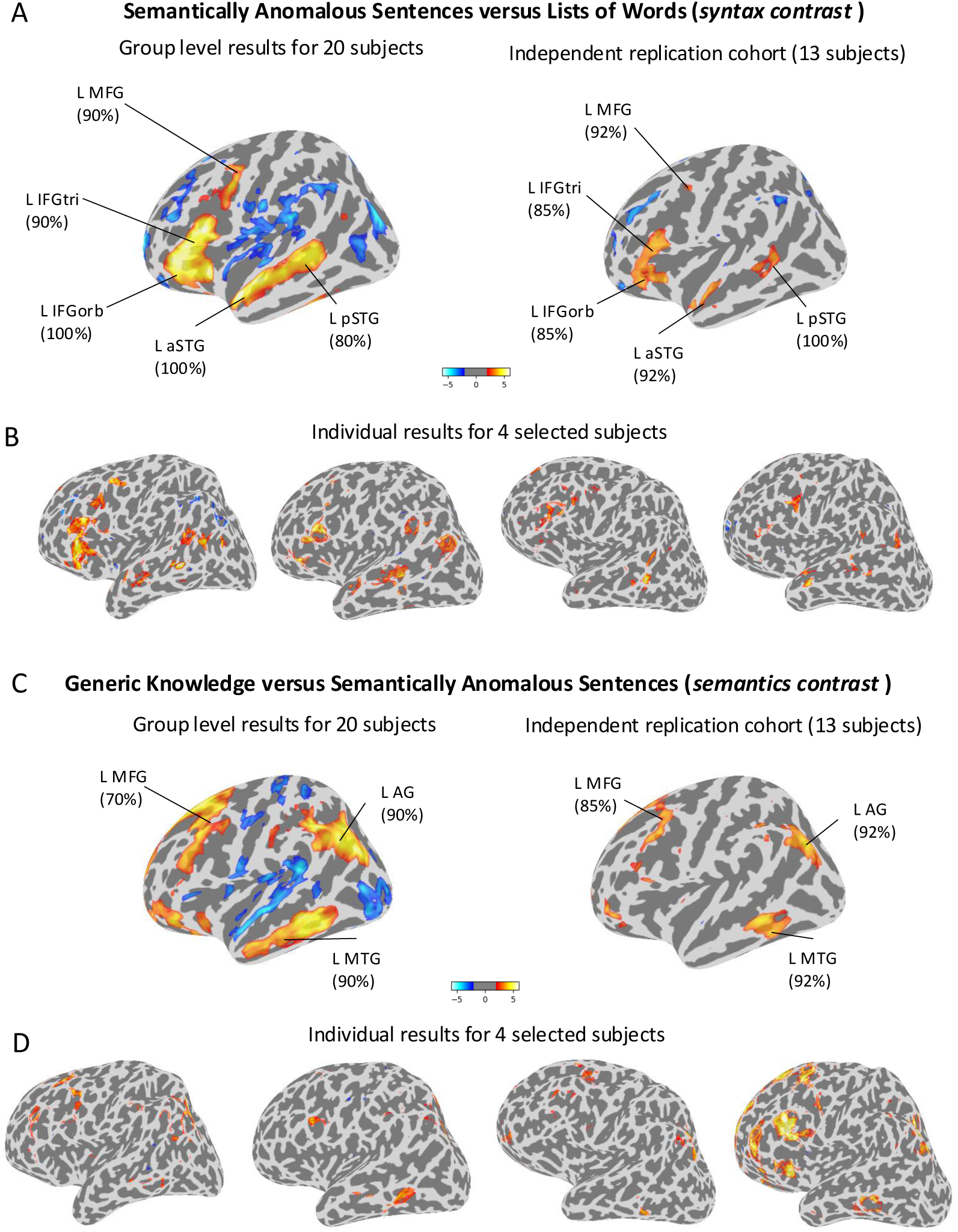
Two distinct networks for processing syntax and semantics. High-resolution 7T fMRI reveals two spatially distinct cortical networks specialized for the extraction of syntactic structure and propositional meaning. A: Group-level contrast between semantically anomalous sentences and lists of words. The contrast between semantically anomalous sentences ("Colorless") and unstructured word lists isolates the core syntactic network. The top row shows the main cohort (n=20); the middle row demonstrates the exact replication of this topography in the independent iCortex cohort (n=13). Results are thresholded at p<0.05, voxel-wise False Discovery Rate (FDR) corrected. Activation is consistently localized to the left superior temporal sulcus (STS) and the inferior frontal gyrus (IFG). B: Individual-level granularity of syntactic patches. Representative activation maps from four individual subjects (thresholded at p<0.001, uncorrected, cluster-extent threshold of k≥4 voxels) reveal the fine-grained distribution of syntactic specialized patches. Key hubs include the left anterior and posterior superior temporal gyrus (L aSTG, L pSTG), middle frontal gyrus (L MFG), and the *pars triangularis* and *orbitalis* of the inferior frontal gyrus (L IFGtri, L IFGorb). C: Group-level contrast between generic knowledge sentences and semantically anomalous sentences. The contrast between factual sentences (generic knowledge) and semantically anomalous sentences ("Colorless") isolates regions involved in truth-value and propositional semantic processing. Top and middle rows show high consistency across the main (n=20) and iCortex (n=13) cohorts (p<0.05, FDR corrected). The network primarily recruits middle and inferior temporal regions, as well as the angular gyrus, largely bypassing the superior temporal syntactic core. D: Individual-level semantic topography. Individual subject maps (thresholded at p<0.001, uncorrected, cluster-extent threshold of k≥4 voxels) illustrate the spatial stability of semantic activation across the left middle temporal gyrus (L MTG), angular gyrus (L AG), and middle frontal gyrus (L MFG). Comparison with panel B suggests that even in native space, these networks occupy non-overlapping cortical territories.

Moving up the linguistic hierarchy, we next examined the activation evoked by generic meaningful sentences relative to semantically anomalous sentences of the same syntactic complexity (referred to as “semantic voxels” for simplicity). At the group level (Figure 2C), this contrast revealed an "extended language network" distinct from the core syntactic regions. In the primary cohort, this semantic contrast primarily recruited the middle temporal gyrus (MTG; 18/20, 90%), the angular gyrus (AG; 18/20, 90%), and the MFG (14/20, 70%). Replication in the second cohort confirmed these findings (MTG: 92%; AG: 92%; MFG: 85%). Individual-subject maps are presented in figure 2D and further detailed in Figures S1–S4. Collectively, these results demonstrate that while syntactic structures evoke a consistent response within the core perisylvian and frontal regions, the addition of semantic content does not just lead to more activity in the same sites, but recruits a second cortical network of surrounding areas, similar to previous findings at 3T (Moreno et al., 2025).

### High-Resolution Individual Analysis Reveals Spatially Segregated Functional Patches

To exploit the enhanced spatial resolution of 7T fMRI, we characterized the anatomical distribution of syntax and semantics patches within each participant’s native space. For this quantitative overlap analysis, we focused on the iCortex replication cohort (n=13), as these participants provided twice the data volume per subject compared to the initial cohort, ensuring a higher signal-to-noise ratio (SNR) necessary for reliable individual-level mapping, and sufficient data to plot the activations in a cross-validated manner.

Quantitative assessment using the Dice Similarity Coefficient (DSC) confirmed a striking degree of spatial segregation between the two networks. The group mean DSC was 0.034 ± 0.033 (mean ± SD), indicating that less than 4% of the active voxels were shared between syntactic and semantic contrasts. Where minimal overlap was observed, it was localized primarily to the anterior temporal lobe (ATL) or the angular gyrus.

This fine-grained analysis unraveled a complex interleaved organization that is often obscured in group-level analyses. Along the superior temporal sulcus (STS), we observed neighboring but distinct patches that maintained a consistent preference for either syntactic or semantic content. Figure 3 illustrates these patterns in three representative subjects: Participant 1 (top) displays an interleaved mosaic of neighboring specialized patches; Participant 2 (middle) shows a distribution more closely aligned with the group-level hierarchy, with STS showing primarily syntactic patches, and MTG primarily semantic patches; and Participant 3 (bottom) exhibits numerous, highly segregated patches with little to no overlap. These individual-level architectures, extended to all 13 iCortex participants in Figure S5, suggest that while both functional systems reside within the same macro-anatomical regions, they occupy distinct cortical territories. In the following, we further explored this coexistence using the multi-modal parcellation proposed by Glasser and colleagues (Glasser et al., 2016).

**Figure 3:**
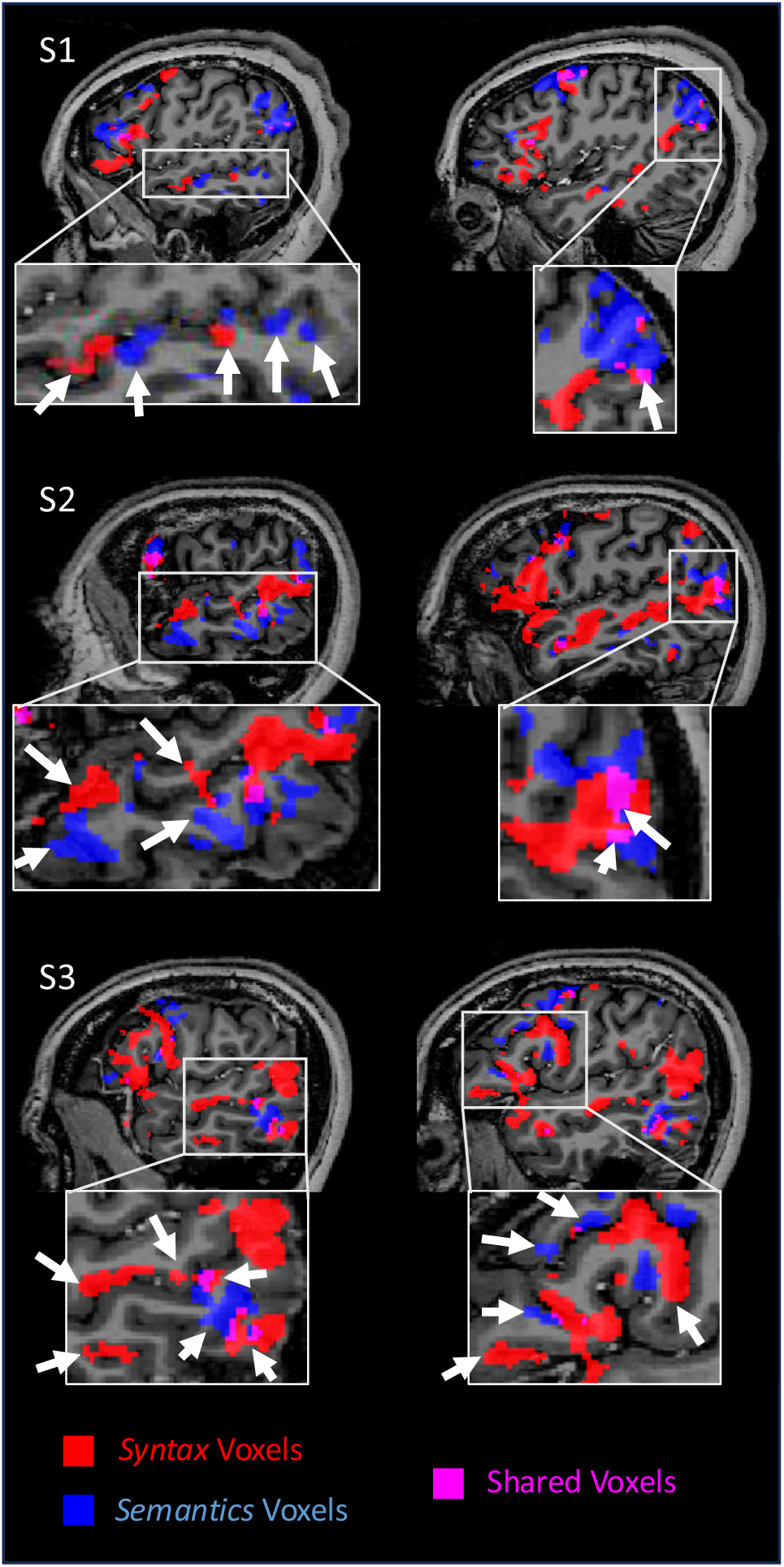
Fine-grained topographical segregation and interleaved organization of syntax and semantics. Single-subject analysis reveals a precise spatial dissociation between syntactic and semantic processing streams within the left temporal lobe. Individual-level cortical mapping: Synthetic activation maps for three representative participants are superimposed on their native T1-weighted anatomy, illustrating the spatial distribution of syntax-responsive (red) and semantics-responsive (blue) voxels. Interleaved functional architecture: Magnified sagittal sections of the left hemisphere demonstrate an "interleaved" organization, particularly along the superior temporal sulcus (STS), where specialized patches for syntax and semantics neighbor one another with minimal spatial overlap (pink). Statistical thresholding: Individual contrast maps are thresholded at p<0.001 (uncorrected) with a cluster-extent threshold of k≥4 voxels to preserve the high-frequency spatial details afforded by the 7T signal.

### Functional decomposition within language-related areas of the Glasser Atlas

The observation of co-occurring activation patches at the individual level suggests that even a single anatomical region may comprise multiple heterogeneous functional voxel populations. To quantify this heterogeneity, we assessed the relative proportions of syntax-related, semantics-related, and shared (overlapping) voxels within each of the 25 language-related regions of the Glasser atlas (specifically HCP-MMP1.0, which comprises a total of 180 relatively small functionally and anatomically defined cortical regions per hemisphere).

As a rule of thumb and to facilitate the comprehension of the results, regions were categorized as syntax-dominant, semantics-dominant or balanced based on a 60% volume-proportion threshold: an area was classified as syntax- or semantics-dominant if its respective specific voxel population exceeded this threshold; otherwise, the region was classified as functionally balanced. Across the 25 regions analyzed, we identified 17 syntax-dominant, 3 semantics-dominant, and 5 functionally balanced regions (see Table 1). Syntactic dominance was most pronounced along the superior temporal sulcus (areas STS_da, STS_dp, and TPOJ1) and within the *pars orbitalis* of the inferior frontal gyrus (areas 45, 47s, and 47l). These findings indicate that while most regions within the language network exhibit a high degree of functional heterogeneity, a clear bias toward syntactic processing exists in core perisylvian and inferior frontal areas.

**Table 1.** Language-related areas of the Glasser atlas differ in their proportion of syntax and semantic-related voxels. For each region, the first two columns report the proportion of voxels with significant activation in the syntax contrast, and in the semantic contrast (p<.001). Parentheses give the number of participants with at least one cluster of 4 significant voxels. The last column gives a tentative label of the profile of the region as being syntax-dominant (Syn-dom, >60% of Syntax-related voxels), semantic-dominant (Sem-dom, >60% of Semantic-related voxels) or balanced.

| **Region** | **% syntax**<br>**voxels** | **% semantic**<br>**voxels** | **% shared**<br>**voxels** | **# of subjects**<br>**(syn contrast /**<br>**sem contrast)** | **Profile** |
| --- | --- | --- | --- | --- | --- |
| **9p_L** | 30.8% | 67.9% | 1.2% | 23 / 27 | Sem-dom |
| **8BL_L** | 26.1% | 72.5% | 1.5% | 24 / 24 | Sem-dom |
| **10pp_L** | 67.0% | 33.0% | 0% | 11 / 5 | Syn-dom |
| **TE1a_L** | 60% | 40% | 0% | 28 / 26 | Balanced |
| **9a_L** | 47.4% | 51.9% | 0.7% | 21 / 26 | Balanced |
| **8Av_L** | 35.0% | 63.8% | 1.18% | 27 / 29 | Sem-dom |
| **TGd_L** | 91% | 8.51% | 0.49% | 18 / 10 | Syn-dom |
| **10v_L** | 52.5% | 47.5% | 0% | 17 / 17 | Balanced |
| **9m_L** | 57.8% | 39.4% | 2.81% | 28 / 28 | Balanced |
| **SFL_L** | 62.8% | 36.1% | 1.12% | 30 / 23 | Syn-dom |
| **PGi_L** | 55.6% | 42.8% | 1.61% | 29 / 28 | Balanced |
| **47s_L** | 88.5% | 11.5% | 0% | 22 / 11 | Syn-dom |
| **44_L** | 74.6% | 24.8% | 0.67% | 27 / 19 | Syn-dom |
| **55b_L** | 73.3% | 25.9% | 0.86% | 26 / 14 | Syn-dom |
| **STSvp_L** | 80.6% | 18.8% | 0.65% | 30 / 18 | Syn-dom |
| **PSL_L** | 70.4% | 29.6% | 0.06% | 22 / 14 | Syn-dom |
| **STSva_L** | 85.6% | 13.3% | 1.04% | 29 / 16 | Syn-dom |
| **47l_L** | 95.4% | 4.57% | 0.03% | 30 / 16 | Syn-dom |
| **STV_L** | 93.8% | 5.58% | 0.60% | 23 / 10 | Syn-dom |
| **45_L** | 92.1% | 7.32% | 0.56% | 31 / 14 | Syn-dom |
| **STGa_L** | 99.7% | 0.29% | 0% | 32 / 1 | Syn-dom |
| **A5_L** | 97.7% | 2.31% | 0% | 30 / 7 | Syn-dom |
| **STSdp_L** | 90.0% | 9.94% | 0.04% | 30 / 10 | Syn-dom |
| **STSda_L** | 98.2% | 1.70% | 0.01% | 30 / 9 | Syn-dom |
| **TPOJ1_L** | 99.3% | 0.74% | 0% | 26 / 4 | Syn-dom |

### Cross-validation of activation profiles to sentences in experiment 1

To evaluate the reproducibility and functional specificity of the identified cortical patches within a Glasser region, we performed an independent cross-validation analysis on the iCortex cohort (n=13). Since participants were exposed to five fMRI runs, for a total of 40 sentences per condition, we could use a 5-fold leave-one-run-out procedure to (1) identify syntactic and semantic voxels from 4 of the 5 runs, and (2) plot the unbiased activation profiles in the fifth run, which used entirely distinct stimuli; (3) rotating across the five runs and averaging the results within-subject; (4) analyzing the results using Linear Mixed-Effects (LME) models and testing for region by condition interactions, a crucial step as stressed by Fedorenko et al. (2024).

The resulting cross-validated profiles are shown in Figure 4 for all language regions averaged together as well as for four typical ROIs (see Figures S7–S8 for all ROIs). Even within a small region such as left BA44, quite distinct voxels were found. In syntax voxels, the cross-validated activation to semantically anomalous sentences was much higher than to lists of words, and reached the same level as meaningful sentences, consistent with the hypothesis that syntactic structure alone suffices to activate them. Conversely, in semantics voxels, semantically anomalous sentences led to a very low activation, often on a par with lists of words, and definitely lower than to meaningful sentences, confirming that these voxels activate in relation to the presence of semantic composition.

**Figure 4:**
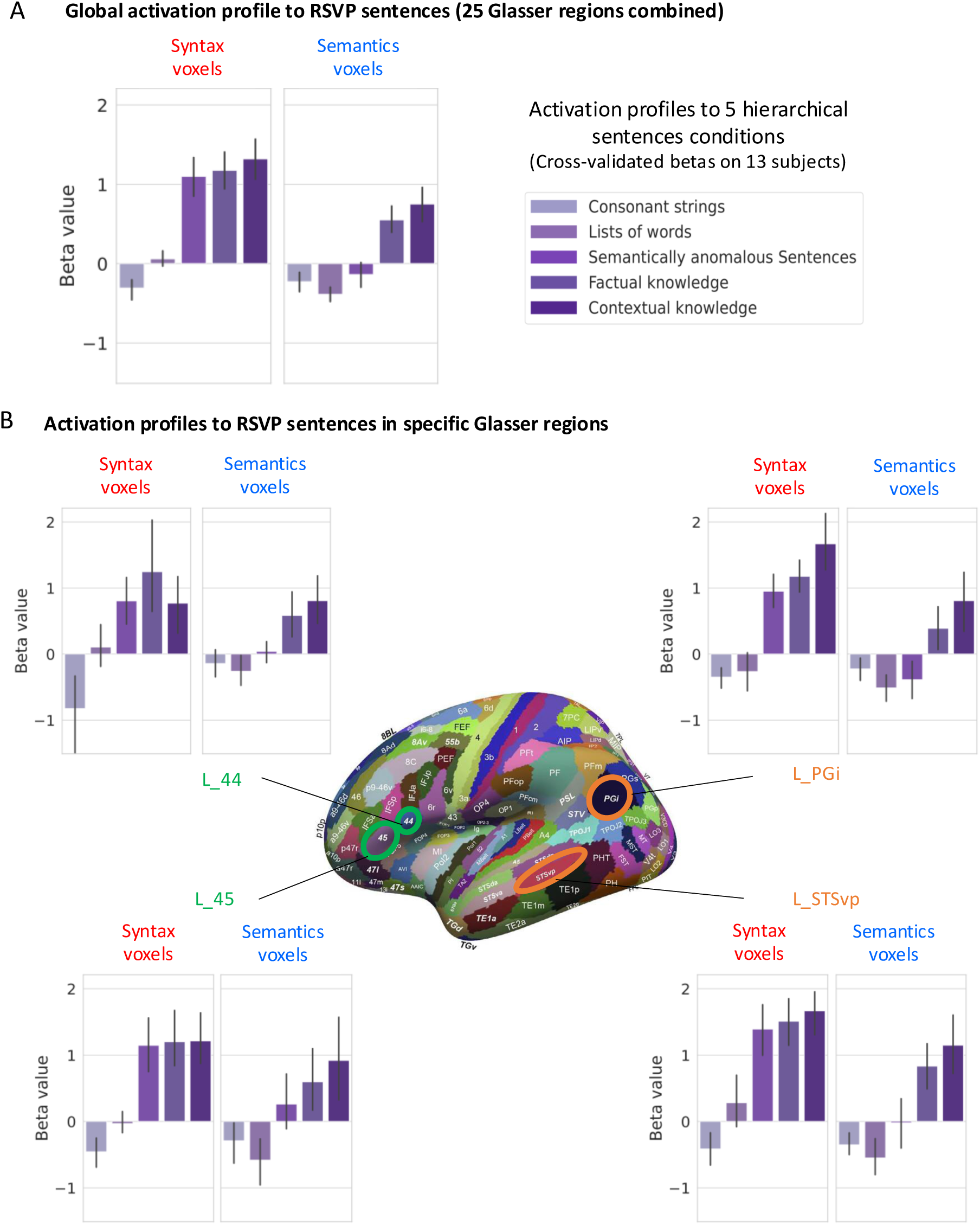
Independent cross-validation of functional profiles within the same anatomical parcels. To confirm the functional specialization and stability of the observed patches, a rigorous leave-one-run-out cross-validation was performed on the iCortex cohort (n=13). Cross-validated voxel selection: Within each subject, five independent runs of Task 1 were analyzed using a split-half procedure. A first-level GLM was trained on four runs to identify syntax-selective and semantics-selective voxels (p<0.001, k≥4 voxels), and their activation profiles were extracted from the independent, left-out fifth run. This procedure was iterated five times, ensuring that the reported beta values are unbiased by the selection criteria. Dissociable profiles within the Glasser atlas: Functional voxels were localized using the multi-modal parcellation of Glasser et al. (2016). Crucially, even within the same anatomical boundaries—such as areas 44 and 45 (Broca’s area), STSvp, and PGi (Angular Gyrus)—two distinct populations of voxels were identified with sharply contrasting response profiles. Functional specialization: In all selected regions, syntax-selective voxels (left graphs) demonstrate a specific sensitivity to syntactic structure, while semantics-selective voxels (right graphs) respond exclusively to the presence of propositional meaning. This within-area heterogeneity underscores the necessity of high-resolution 7T imaging to resolve the fine-grained functional architecture of the language network.

Activation profiles were analyzed using Linear Mixed-Effects (LME) models, with a random slope for voxel type per subject fit by classical weighted least squares with weights equal to n_voxels_ to reflect the precision of per-subject beta means. Tests for voxel-type × condition interactions were performed at two levels: global, and within each of the 25 language regions.

Prior to any regional decomposition, we tested whether the voxel-type × condition interaction was detectable in a pooled model that did not include region as a covariate — that is, treating all syntax and semantic voxels across the language network as a single pool. This model revealed a highly significant interaction (*χ*^2^(4) = 630.88, *p* < 10^−168^), confirming that the two voxel populations exhibit fundamentally different condition profiles even without accounting for anatomical location.

A global LME statistical analysis across all 25 language regions, including regions as a fixed effect, revealed a highly significant voxel-type × condition interaction (*χ*^2^(4) = 785.14, *p* < 10^−168^), confirming that syntax and semantics patches exhibit fundamentally different responses across the linguistic hierarchy. Crucially, we quantified heterogeneity of this interaction across regions — the interpretable form of a region × voxel-type × condition triple interaction on unbalanced data — using Cochran’s Q on the per-region z-transformed p-values, weighted by the number of dual-patch subjects, i.e., the number of participants with both type of voxels in a given region. This heterogeneity was highly significant (*Q*(24) = 259.0, *p* < 10⁻¹⁶), statistically demonstrating that the functional dissociation varied across anatomical regions. Sensitivity analyses of the LME under unweighted and square-root weighting variants showed the same pattern across regions (Supplementary Tables S1, S1b).

LME analyses restricted to each Glasser area confirmed significant Voxel-type x Condition interactions in 18 of the 25 testable language regions after FDR correction (Table 2). A primary driver of these effects was the response to semantically anomalous sentences, to which syntax-tuned voxels maintained high activation levels (close to, though slightly lower than, meaningful sentences), whereas semantic-tuned voxels showed a sharp reduction in BOLD signal. This dissociation was found in most of the tested regions. In semantic voxels, we found reduced activation for semantically anomalous sentences compared to meaningful sentences in 23 of 25 regions after FDR correction. In syntax voxels only 5 of 25 regions showed a similar pattern of increased activation to meaningful sentences compared to semantically anomalous sentences (10v_L, 8BL_L, 9m_L, PGi_L, TGd_L), while 4 regions showed the opposite pattern (47l_L, A5_L, STSda_L, TPOJ1_L) and the remaining 16 regions showed no significant differences between both conditions.

**Table 2.**
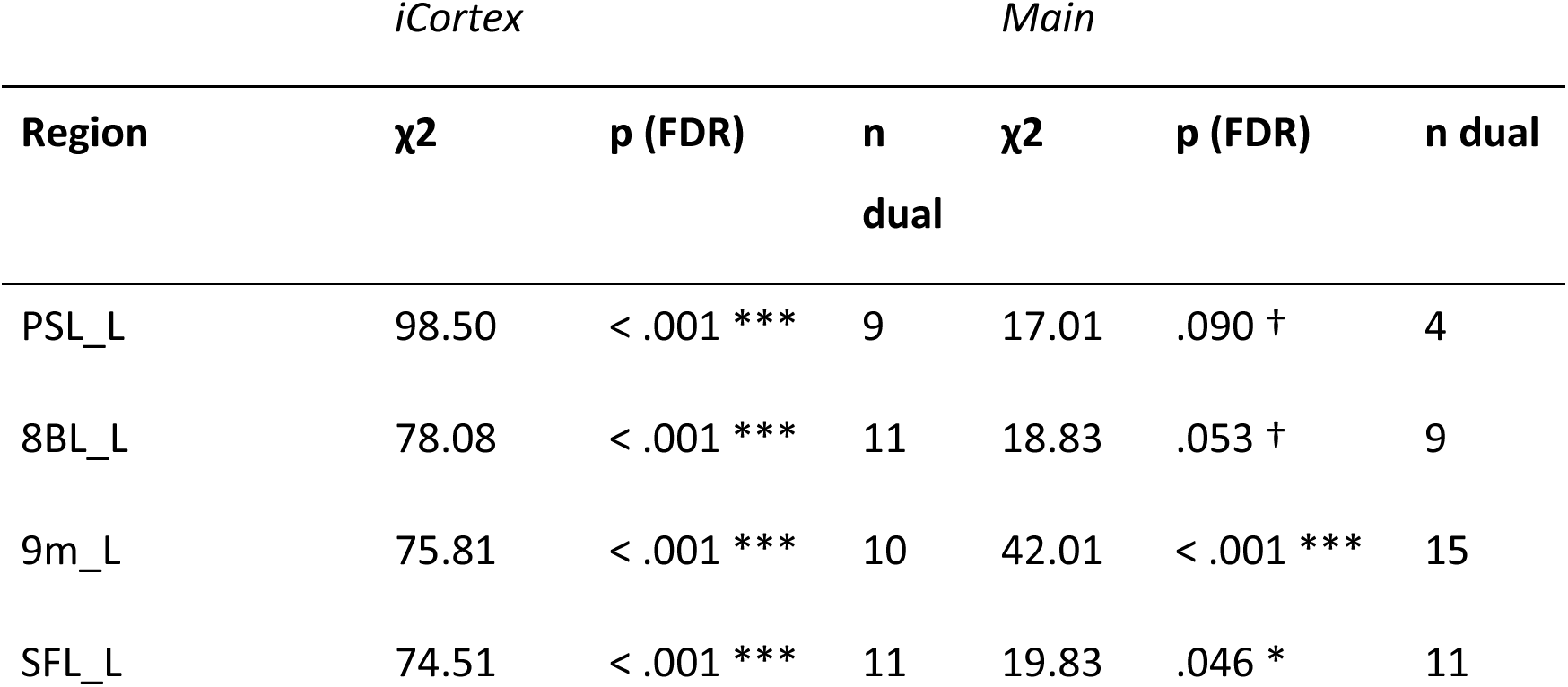
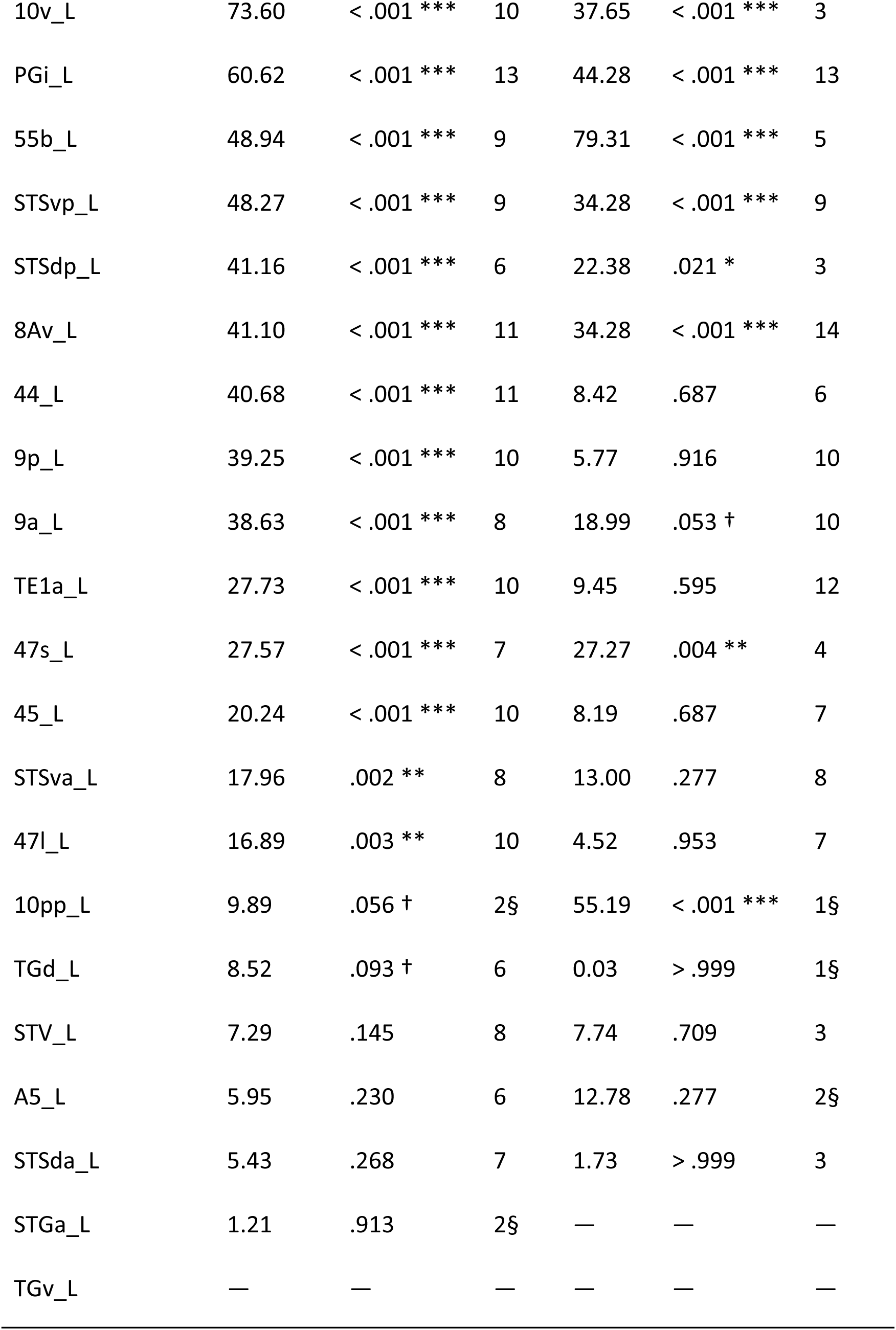
Statistical tests evaluating whether, within each Glasser area, syntax- and semantic-related voxels differ in their activation profiles to linguistic stimuli. For each Glasser area, voxels were separated into those responsive to syntax versus semantic contrasts, and then tested for a condition X voxel type interaction using a mixed-effect linear model on an independent data set. Left, test for differences between the 5 conditions of the RSVP truth judgement task (see Figure 4 for a plot of those activation profiles); right, test for differences between the 10 mini-sentences conditions (see Figure 6 for a plot of those activation profiles). *Note.* For each region, voxels were separated into syntax- and semantic-defining populations, and a voxel-type × condition interaction was tested using a Linear Mixed-Effects model with a random slope for voxel type per subject, fit by classical weighted least squares with weights = *n*voxels (linear voxel-count weighting). The *iCortex* columns report results from the RSVP task (5 conditions, *n* = 13); the *Main* columns report results from the mini-sentences task (10 conditions, *n* = 20). *n* dual = number of participants with at least one voxel of each type in the region. **§** indicates cells where *n* dual ≤ 2: in these regions the interaction is largely driven by between-subject variation rather than within-subject dissociation, because fewer than three subjects carry both voxel types. These rows should be interpreted as sample-level effects only. Em-dashes indicate cells where the model could not be fit (no voxels of one type, or fewer than two distinct subjects). Sensitivity analyses under unweighted and square-root weighting are in Supplementary Tables S1 and S1b. † *p* < .10. * *p* < .05. ** *p* < .01. *** *p* < .001.

Systematic comparison between the activation to semantically anomalous sentences and to lists of words, using the same weighted paired-*t* approach with FDR correction within each voxel type, confirmed the dissociation. Syntax voxels showed significantly greater activation to anomalous sentences than to word lists in 24 of 25 regions after FDR correction, confirming that well-structured sentences — even when semantically deviant — drive syntax voxels substantially more than lists of words. For semantic voxels, the same contrast was significant in only 4 of 25 regions (9m_L, STSva_L, STV_L, TE1a_L; the 21 other regions were non-significant, as no region showed a significant negative difference). This dissociation is consistent with the interpretation that syntactic processing — not semantic content — is the primary driver of the differential response in syntax voxels between sentences devoid of overall meaning and lists of words without syntactic structure.

We verified that the observed dissociation was not driven by the extreme conditions (namely lists of consonant strings and contextual sentences). When restricting analyses to the 3 central conditions (word lists, semantically anomalous sentences, factual knowledge sentences), the pooled no-region model on this three-condition subset remained highly significant (*χ*^2^(2) = 300.28, *p* < 10^−64^), and the region-inclusive variant likewise (*χ*^2^(2) = 429.10, *p* < 10^−93^). Per-region tests on the restricted subset yielded significant voxel-type × condition interactions in 19 of the 25 testable regions after FDR correction, compared with 18 of 25 in the full five-condition model. The consistency of results within this subset confirms that the dissociation is robustly driven by the three core linguistic conditions and does not depend on the inclusion of the pseudoword or contextual knowledge conditions.

### Behavioral and Neural Responses to Minimal Syntactic Structure

Experiment 2 aimed to examine the neural sensitivity of the same voxels to minimal syntactic operations, thus checking that they indeed implement distinct operations. On each trial, participant were flashed a single letter string designed to minimize semantic load while varying structural complexity. The stimuli spanned 10 categories including three control conditions (single words, word lists, and ungrammatical strings) and seven grammatical conditions involving basic sentences (e.g. “you go there”, “I see it”) as well as minimal syntactic manipulations (e.g. wh-questions, etc; Fig. 1B).

During fMRI, participants merely read the mini-sentences passively, waiting for a rare cue to perform a match-to-sample task. To verify that the flashed stimuli could be perceived and parsed, and that our manipulation of syntactic complexity worked, we also collected behavioral performance on a distinct post-scanning syntactic decision task, conducted with the same 200 ms Rapid Parallel Visual Presentation (RPVP), where participants decided whether the stimulus was a well-formed sentence in French. Behavioral results confirmed that syntactic complexity significantly modulated processing time (Fig. 5). Error rates and median reaction times (RTs) were highest for ungrammatical sentences, where word order violated French syntactic constraints: ungrammatical sentences resulted in significantly higher error rates (Mean: 15.9%) and median reaction times (Mean: 607 ms) compared to grammatical ones (3.5%; 491 ms). We included these results in a repeated measures ANOVA and found a highly significant effect of syntactic complexity on reaction times (F_1,19_ = 56.5, p<0.001) and error rates (F_1,19_ =47.3, p<0.001). Within the grammatical categories, increasing syntactic complexity (e.g. due to presence of clitic movement or verb movement in wh-questions) resulted in a systematic increase in both RTs (F_2,38_ =26.1, p<0.001) and error rates (F_2,38_ =15.5, p<0.001) (Fig. 5). RTs increased significantly when moving from "0 movement" (463 ms; average of simple affirmative sentences, yes-no questions and wh-in situ questions) to conditions with movement (522 ms = average of 1-movement conditions, i.e., clitic movement, yes-no with verb movement and pure wh-movement; 519 ms = response time to wh plus verb movement).

**Figure 5:**
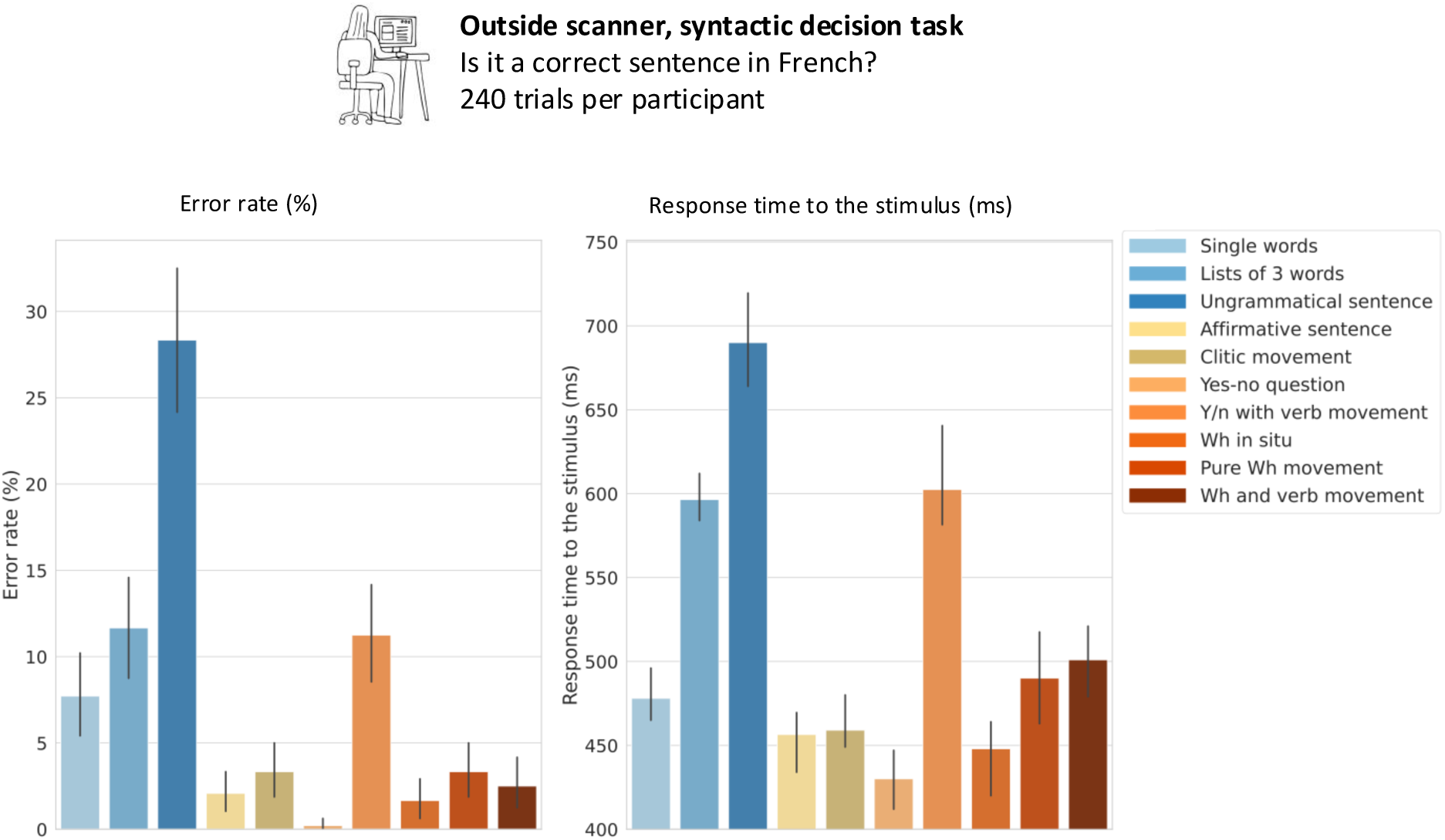
Behavioral signatures of minimal syntactic complexity. To validate the hierarchical structure of the "single-glance" stimuli, behavioral performance was assessed using a syntactic decision task matching the 200 ms Rapid Parallel Visual Presentation (RPVP) timing used during fMRI. Complexity-dependent processing costs: Both error rates (left) and median reaction times (right) exhibit a systematic increase as a function of syntactic complexity. Sensitivity to grammatical constraints: Processing costs are maximized for ungrammatical strings that violate French syntactic constraints, reflecting a significant "grammaticality effect." Validation of minimal structure: Within the grammatical categories, the transition from simple structures to those involving clitic, verb or wh-movement (conditions 5, 7, 9 and 10) results in a linear increase in difficulty. High-speed robustness: These results confirm that the language system is capable of extracting complex structural information even under the stringent temporal constraints of a 200 ms presentation.

These out-of-fMRI behavioral results validate the hierarchical nature of the stimuli and confirm that participants were sensitive to the underlying structural manipulations even at high-speed visual presentation, suggesting that the same effects must have occurred covertly during the scanning (where we purposedly did not collect responses on the analyzed trials, to avoid task-and decision-related confounds). Behavioral results of the match-to-sample task during fMRI did not reveal a sensitivity to syntactic complexity, instead suggesting that all stimuli were roughly equally well remembered (Fig. S6). We performed the same repeated measures ANOVA. There was no significant difference in the time subjects took to respond to ungrammatical sentences (958 ms) compared to grammatical ones (966 ms) (F1,19 =0.007,p=0.934.) or in the error rates (Error Rate: 10.2% for Grammatical vs 9.0% for Ungrammatical) (F1,19 =0.356,p=0.558). Syntactic complexity did not affect RTs (F1,19 =2.283,p=0.147) nor error rates (F1,19 =1.035,p=0.322).

### fMRI responses to mini-sentences confirm the dissociation of functional subnetworks

We then examined how the subject-specific voxels identified as syntax- or semantics-related in experiment 1 responded to the 10 mini-sentence conditions (Figures 6, S10–S11). We first tested whether the voxel-type × condition interaction was detectable across all language regions in a single pooled model that excluded region as a covariate, treating all syntax and semantic voxels as a single population. This pooled LME revealed a highly significant interaction (*χ*^2^(9) = 92.20, *p* = 5.6 × 10^−16^), confirming that the two voxel types exhibit qualitatively different response profiles across the 10 mini-sentence conditions even before accounting for regional structure.

**Figure 6:**
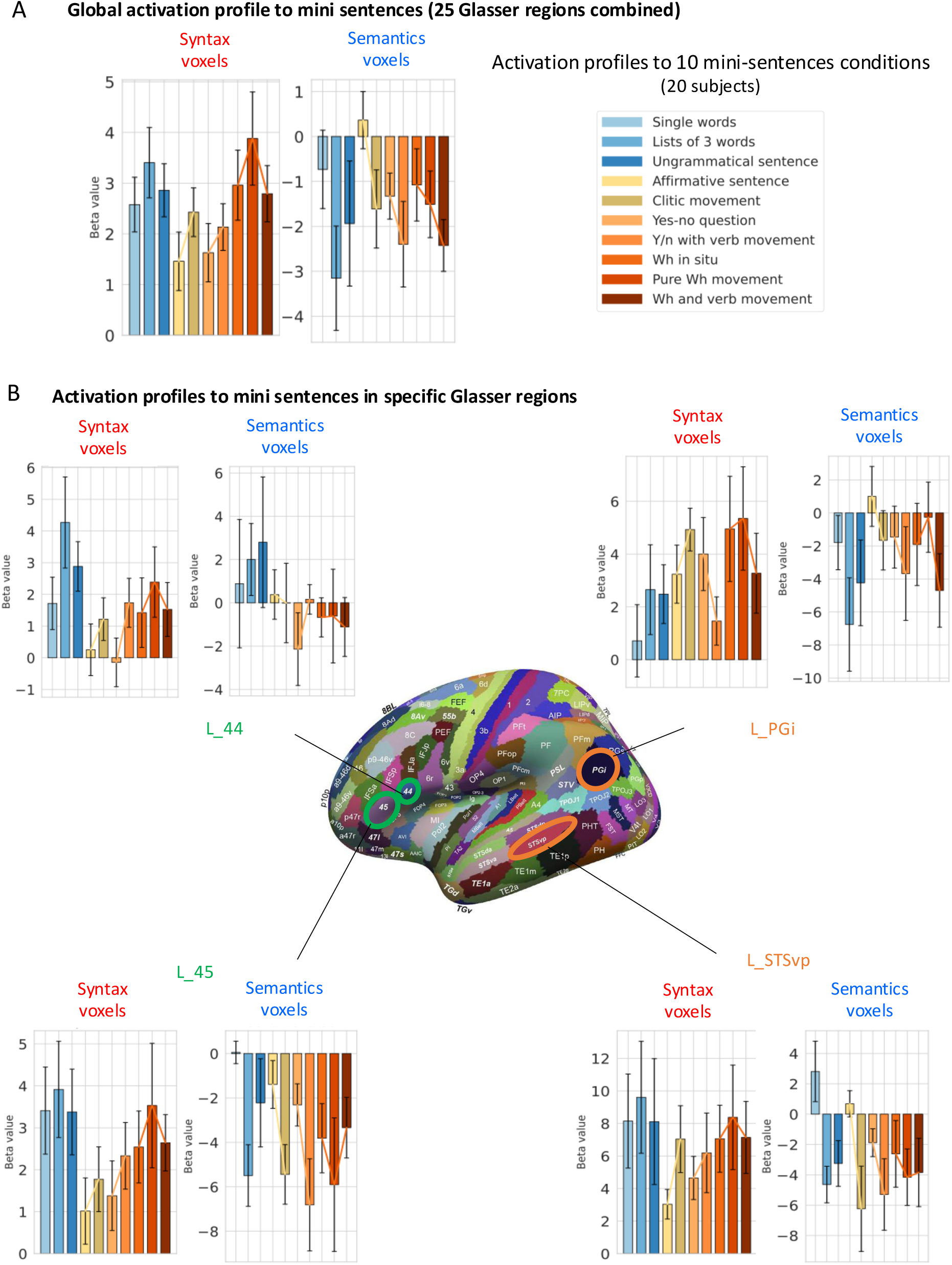
Generalization of syntax- and semantics-selective profiles to minimal linguistic structures. To test the robustness of the identified functional patches, we examined whether voxels defined by the RSVP truth judgement task (Task 1) maintain their specialized response profiles when processing "single-glance" minimal sentences (Task 2). Independent functional localization: For each participant, syntax-selective and semantics-selective voxels were independently localized using the contrasts from Task 1 (Semantically anomalous vs. Word lists and Factual vs. Semantically anomalous, respectively). Selection was performed at the individual level (p<0.001, uncorrected) and voxels were assigned to anatomical parcels using the Glasser atlas. Response profiles across the linguistic hierarchy: Beta values were extracted for the ten stimulus categories of Task 2, which range from unstructured controls to complex grammatical constructions. In key hubs—including Broca’s area (L 44, L 45), the superior temporal sulcus (L STSvp), and the angular gyrus (L PGi)—the two voxel populations exhibit sharply divergent sensitivities. Functional invariance across scales: Despite the drastic change in presentation format (RSVP vs. 200 ms single-glance) and complexity (full sentences vs. 3-word strings), the syntax-defined voxels (left) consistently track structural manipulations, with larger activation to more complex stimuli, while the semantics-defined voxels (right) are largely deactivated, with an inverted modulation by syntactic complexity. The stability of the dissociation of Syntax and Semantics voxels across tasks provides strong evidence for a fixed, fine-grained topographical organization of the language network.

We next moved to a region-inclusive global LME where region is included as a fixed-effect covariate. A global LME statistical analysis across all 25 language regions, including regions as a fixed effect, revealed a highly significant voxel-type × condition interaction (*χ*^2^(9) = 105.81, *p* < 10^−18^), with significant heterogeneity of the interaction across regions (Cochran’s *Q*(23) = 546.48, *p* < 10^−16^). Per-region tests were less sensitive on this dataset than on iCortex, reflecting both the larger number of conditions (10 vs 5) and the typically smaller voxel counts obtained in the main cohort, exposed to half as many RSVP sentences as iCortex participants. Nevertheless, per-region tests on each of the 24 testable regions (TGv_L and STGa_L lacked sufficient dual-voxel coverage) yielded 10 of 24 regions significant after FDR correction (Table 2). The significant regions with adequate dual-voxel coverage were 47s_L, 55b_L, 8Av_L, 9m_L, 10v_L, PGi_L, SFL_L, STSdp_L, and STSvp_L (all *p*_FDR ≤ .046); a tenth region (10pp_L, *n*_dual = 1) reached significance but reflects between-subject contributions only (see Table 2 footnote). Three additional regions were trending (8BL_L, 9a_L, PSL_L; all *p*_FDR < .10). Such interactions, using independent data from those used to select the voxels, fill the criteria suggested by Fedorenko et al. (2024) as indicating a genuine functional dissociation.

Examination of the histograms of activation (beta values) to each of the 10 mini-sentence conditions revealed the nature of this dissociation (see Fig. 6, and S10-S11 for the full set of ROIs). Strikingly, in virtually all language areas, syntax patches exhibited a strong activation to single-glance stimuli relative to rest (positive response to the 7 grammatical sentence conditions relative to rest ; all *p*_FDR < .01), whereas semantics patches exhibited near-zero activation or even a deactivation (their activation to the simplest grammatical condition, affirmative sentences, was not significantly different from zero, *p*_FDR = .333, while activation to the six remaining sentence conditions was significantly negative after FDR correction, all *p*_FDR ≤ .05). This finding is likely due to the fact that mini-sentences were syntactically rich but semantically impoverished, since they comprised pronouns and locatives devoid of any semantic referent (e.g. he, it, there).

We next asked whether, within the 7 categories of syntactically well-formed sentences, an effect of syntactic movement was seen. An LME restricted to the grammatical subset again revealed a significant voxel-type × condition interaction (*χ*^2^(6) = 91.77, *p* < 10^−17^), with significant cross-region heterogeneity (*Q*(23) = 276.06, *p* < 10^−16^). To test the directionality of this effect with respect to movement complexity, we coded each grammatical condition as carrying either no syntactic movement (conditions 04, 06, 08) or at least one movement (conditions 05, 07, 09, 10), and fit an LME with movement (0/1) × voxel-type as fixed effects, region as a covariate, and a random slope for voxel type per subject. Remarkably, syntax and semantics voxels responded to the presence of movement in opposite directions. As expected, syntax voxels showed a positive slope, meaning that activation increased with syntactic complexity (β = +0.79, *SE* = 0.18, *z* = +4.39, *p* < 10^−4^). However, remarkably, the opposite was true for semantics voxels: they showed a negative slope, indicating that increased syntactic complexity lead to lesser activations (β = −1.31, *SE* = 0.29, *z* = −4.54, *p* < 10^−5^). Thus, the voxel-type by movement interaction was highly significant (*χ*^2^(1) = 38.16, *p* < 10^−9^). Running the same model without including regions as a covariate yielded nearly identical results: both slopes individually reached significance and pointed in opposite directions, confirming that syntax and semantic voxels responded to the presence of syntactic movement in qualitatively different ways.

Refitting the same model independently in each of the 22 testable regions (TGv_L and STGa_L lacked any semantic voxels) confirmed that this dissociation was not driven by a small subset of regions. The slope in semantic voxels was negative in 18 regions (FDR-significant in 8: 55b_L, 8Av_L, 8BL_L, 9a_L, 9m_L, PGi_L, PSL_L, STSvp_L) and never significantly positive, while the slope for syntax voxels was positive in 13 regions (FDR-significant in 8: 44_L, 45_L, 55b_L, PSL_L, SFL_L, STSda_L, STSvp_L, TPOJ1_L) and significantly negative in only one (9a_L). Crucially, no region exhibited the reverse pattern (positive slope for semantic voxels combined with negative slope for syntax voxels), and 11 regions reproduced the pooled dissociation in point-estimate form. The per-region interaction reached FDR significance in four regions (55b_L, 8Av_L, PSL_L, STSvp_L; all *p*_FDR_ < .002), with three further regions trending (8BL_L, 9m_L, SFL_L; all *p*_FDR_ < .10; Supplementary Table S6). The pooled effect therefore reflects a network-wide property of the language regions rather than an effect localized to one or two patches.

### Sensitivity of the syntax network to structural manipulations

Having established that syntax voxels respond strongly to well-formed sentences relative to lists of words, we asked how their activation was modulated by the type of syntactic structure carried by each sentence. As a whole, the pattern of activation for syntax and semantic voxels across all 25 regions was striking: syntax voxels exhibited a gradual increase in activation from basic affirmative sentences to those involving clitic, verb, or wh movement, and a small reduction when wh and verb movement co-occurred. The latter was previously observed in unpublished work from our lab (Fabre, 2017) and is compatible with theoretical approaches (Rizzi, 1996) which suggest that the wh + verb construction (e.g., où vas tu?) is less marked than wh-movement alone (où tu vas?). The profile was particularly visible in the superior temporal region L_STSvp and the inferior frontal regions L_44 and L_45 (Fig. 6, panel B) and was entirely absent in semantic voxels (Fig. 6, panel A).

To evaluate the role of each specific type of movement, we performed four weighted paired t-tests on syntax voxels in all syntax voxels pooled together and in each of these three ROIs, targeting (1) clitic movement against basic affirmatives (05 − 04), (2) yes-no questions with verb movement against those without (07 − 06), (3) pure wh-movement against wh in situ (09 − 08), and (4) the addition of verb movement to wh-movement (10 − 09). Those individual contrasts were probably underpowered, however, because they all showed a trend but did not reach significance across participants even when syntax voxels were pooled across all language regions (clitic movement 05 − 04: t(19) = 1.72, p = .101; verb in yes-no question 07 − 06: t(19) = 1.17, *p* = .256; pure wh-movement 09 − 08: t(19) = 1.07, *p* = .299; wh + verb vs wh 10 − 09: t(19) = −1.32, *p* = .204; all p_FDR = .299). When the three regions thought to be crucial for syntax, i.e. L_STSvp, L_44 and L_45 (Pallier et al., 2011) were analyzed individually (4 contrasts X 3 regions), only one contrast survived FDR correction (verb movement in L_44, *t*(16) = +3.11, *p*_FDR = .046), with another effect significant at uncorrected level (clitic movement in L_STSvp, *t*(17) = +2.54, *p* = .021, *p*_FDR = .124); the remaining ten contrasts trended in the expected direction without reaching significance. Nevertheless, the directional consistency in the three key ROIs is nonetheless improbable under the null hypothesis (12/12 signed contrasts in the predicted direction, one-tailed sign test, *p* = 2.4 × 10^−4^). Supplementary Table S3 gives the full list of contrasts across all regions.

More surprisingly, syntax-related voxels also activated quite strongly to ungrammatical sentences, often up to the same level as to wh-movement sentences (Fig. 6) and, to a lesser extent, to word lists and long single words. These activations become less surprising, however, when one observes that they parallel the difficulty levels observed in the off-line syntactic decision task (Fig. 5). Since participants were asked to remember the stimuli in order to respond to an occasional one-back probe, the syntactic networks of the brain may have been actively engaged in trying to memorize, parse and/or repair those anomalous stimuli which, as revealed by behavior, were more difficult to process, thus yielding a larger activation. This interpretation was confirmed by a whole-brain group comparison, which indicated that, during the single-glance experiment 2 only, anomalous sentences and word stimuli always yielded significantly *more* activation than basic sentences (analyses presented in supplementary materials and in Fig. S9).

## Discussion

Using 7T functional MRI, our study demonstrates the coexistence of two populations of cortical patches in the language network with different degrees of specialization for syntactic processing and semantic composition. Each participant performed two visual reading experiments. Experiment 1 was designed to functionally localize the language network of each participant and disentangle sub-networks dedicated to semantics and syntax, while Experiment 2 studied the activation profile of the language network to mini-sentences with controlled syntactic manipulations. Even within small language-related regions such as left Brodmann area 44, we indeed found two populations of voxels. The first population activated in response to syntactic structure even in the absence of semantic coherence (i.e. in response to semantically anomalous sentences analogous to Chomsky’s “colorless green ideas…”). The second population showed little or no difference between lists of words and semantically anomalous sentences, but required the sentence to be meaningful in order to attain a strong level of activation. Crucially, this syntax/semantics dissociation was confirmed using cross-validation on Experiment 1 from an independent set of participants, and further backed by the unique profile of activation of both voxel types to the mini-sentences: only syntax-related voxels showed a systematic increase in activation whenever syntactic complexity increased, due either to clitic, to verb, or to wh movement. Semantics-related voxels actually showed further deactivation in response to the same conditions.

### The syntax versus semantics debate

The hypothesis of the existence of syntactic hubs in the language network dates back to the early days of functional brain imaging and has since received a large body of evidence (Bornkessel-Schlesewsky & Schlesewsky, 2013; Dapretto & Bookheimer, 1999; Duffau et al., 2014; Frankland & Greene, 2015; Friederici, 2003; Hagoort, 2005, 2013; Hickok & Poeppel, 2007; Nelson et al., 2017; Tyler et al., 2008, 2010, 2011). Most studies point to key syntax nodes in the posterior temporal and inferior frontal regions, although the exact anatomical localization may slightly differ, partly due to differences in stimuli and experimental design. For instance, in an fMRI study, Tyler et al. (2010) found these regions to be selectively activated by syntactically coherent sentences without meaning. In another study the following year, Tyler et al. (2011) obtained similar results on a cohort of patients recovering from left-hemispheric strokes listening to sentences with syntactic ambiguity. Goucha and Friederici (2015) found activations restricted to pSTG and IFG (Broadman areas 44 and 45) in Jabberwocky conditions as opposed to random word order. More minimalistic paradigms, using two-word or single-word stimuli sufficed to activate the same putative syntactic hubs (Longe et al., 2007; Tyler et al., 2008). The causal role of these syntactic hubs in processing syntactic operations was further supported by a Transcranial direct current stimulation (t-DCS) study by Krause and colleagues (Krause et al., 2023).

The results of the present study also align nicely with those from Pallier, Devauchelle and Dehaene (2011), who identified the posterior superior temporal gyrus (pSTG) and the inferior frontal gyrus (IFG) as putative syntax-related regions on the basis of their showing parallel profiles of increasing activation to constituent size in both Jabberwocky and regular sentences, suggesting that syntax suffice to drive them. Other regions, for instance in the anterior STG, only showed such an increase in response to regular meaningful sentences, and thus were thought to be involved in semantic composition. A similar complexity effect was also observed in French Sign Language and involved the same inferior frontal and temporal regions of the left hemisphere (Moreno et al., 2018). The same set of regions were also involved in the manipulation of syntactic trees (Pattamadilok et al., 2016).

More directly related to our study are previous attempts to identify brain areas involved in syntactic movement. Syntactic movement is a form of the merge operation, the building block of modern minimalist linguistic theory, defined as internal merge (Rizzi, 2012). In a classical study conducted in Hebrew, Ben-Shachar et al. (2004) identified regions in the IFG and posterior STG that showed activation in the presence of syntactic wh movement, namely topicalization and Wh-movement. These results were replicated with a whole new set of sentences and a different task, with greater focus on distinguishing between wh-movement and verb-movement (Shetreet & Friedmann, 2014). These results were extended to other languages such as Japanese (Ueno & Kluender, 2009) and Korean (Park et al., 2024). To this background, our study both behavioral and brain-imaging that syntactic movement induces a significant cost, by systematically increasing activation in syntax but not semantics voxels. Rather than being restricted to specific regions, the present evidence suggests that movement operations involve a distributed activation across syntax-related populations in most if not all language areas.

Surprisingly, in Experiment 2, the contrast between declarative sentences and lists of words or ungrammatical strings identified voxels in the same regions (pSTG and IFG), but now these were more robustly activated by ungrammatical stimuli. We interpret this finding as increased computations in these syntactic hubs in trying to resolve a proper syntactic structure, before reaching the conclusion that the stimuli are indeed ungrammatical. It is important to note that the task was different in Experiments 1 and 2: during sentence verification (Experiment 1), the syntactically anomalous stimuli such as lists of words could be quickly rejected, thus requiring less work than syntactically or semantically organized stimuli, whereas those stimuli were actually more difficulty during a memorization (Experiment 2).

A related argument may explain why the semantically anomalous sentences in Experiment 1 caused a non-zero activation, relative to word lists, even in semantic voxels. Deciding that a sentence makes no sense is not immediate. The first few words of semantically anomalous sentences always made sense, and although we endeavored to make sure that semantic violations were maximal and came as early as possible in the sentence, it is likely that semantic integration started and then failed, thus accounting for the small semantic activations observed. Future work should endeavor to test this hypothesis using time-resolved, word-by-word methods.

An intriguing new finding of our study is the inverse effect of syntactic complexity on the activation profiles of syntax- and semantics-related voxels. In syntax voxels, the observed increasing activation in response to greater syntactic movement was fully expected and consistent with an increased computational load for structural processing. However, semantics voxels exhibited a significant decrease in activation (or even increased deactivation) in response to our mini-sentences. The harder those stimuli were to parse, the more they deactivated the semantic-related voxels. This opposing sensitivity suggests a functional trade-off or a competitive allocation of resources within the language network. It is possible that as the demand for structural reanalysis increases (e.g., in cases of complex movement or topicalization), the cognitive resources available for semantic composition are diminished, or alternatively, that the system deprioritizes semantic integration when structural processing becomes the primary bottleneck. It also seems plausible that this result may be specific to the mini-sentences we used, which involved very minimal possibilities of semantic construction, particularly since they comprise pronouns and locatives without semantic referents.

Our results suggest a tentative reconciliation of the debate between distributed sensitivity and regional specialization in the language domain. While sensitivity to syntactic movement was detectable across a broad range of language-responsive regions, the whole-brain group-level contrasts between declarative sentences and non-sentences (both word lists and ungrammatical strings) identified a discrete set of highly specialized hotspots. These regions, located primarily in the pSTG and IFG, appear to house a significantly higher density of syntax-related voxels. Consequently, our data suggests a "center-periphery" organization: while syntactic information may be processed in a distributed manner throughout the language network, there remain specialized hubs where the neural code for syntax is most concentrated and robust. This explains why broad whole-brain contrasts reliably isolate these specific regions, even if more sensitive, individual-level analyses reveal a wider distribution of syntactic sensitivity.

### Methodological Factors that may explain contradictory findings

Although backed by a rich body of evidence, the existence of regions of the language network dedicated to syntax has been questioned by recent work (Fedorenko et al., 2020, 2024; Shain et al., 2024). Authors of these studies advocate for a distributed sensitivity to both syntax and semantics across the language network rather than specific syntactic hubs. There are, however, several methodological factors that might account for the differences between these studies and ours. First, while presenting the great advantage of focusing on single-subjects, without group-level smoothing, those studies typically identify voxels involved in the language network by contrasting meaningful sentences with lists of pseudo-words, and then pooling together all the voxels that respond to this contrast, typically within a relatively large brain region (Fedorenko et al., 2010). As a result, they run the risk of pooling together voxels with quite different functional properties. As demonstrated here, within subjects, syntax- and semantics-related patches can be intermingled, with the semantic network often having a smaller cortical footprint across most language-responsive ROIs – and while they differ in how they respond to syntactically correct but semantically anomalous sentences, they all show the basic difference between sentences and word or pseudoword lists, and hence would have been pooled together by Fedorenko’s method. Such pulling, using an exceedingly broad language contrast, may fail to isolate functionally specific voxels and create the impression that all regions perform the same computations.

Second, these studies rely on 5 classical yet large fronto-temporal language ROIs. While these regions are especially relevant in group analysis, they lack the precision needed to identify subpopulations of voxels in the language network at the individual level. Indeed, our single-subject analysis shows that even though each participant exhibits distinct syntax and semantics patches, their anatomical distribution does not strictly respect group-defined ROIs such as the areas of Glasser atlas: within most of these areas, both profiles coexist.

Overall, by using a finer-grained analysis of the language network, our study provides clear evidence for syntactic specialization. The observed single-subject syntactic hubs mostly match the past reports from group studies while depicting functional specialization at 7T with a precision that goes beyond the previously reported classical macroscopic brain networks. Indeed, the individual analysis of our 33 participants, divided in 2 groups (20 + 13), revealed patterns of activation that did not strictly respect the anatomical distinctions. Instead of entire brain areas dedicated to specific functions, our results suggest a more anatomically distributed division between syntax and semantics. We used the anatomical and functional brain parcellation by Glasser and Van Essen to conduct our analysis, which provides a set of 180 areas per brain hemisphere (Glasser et al., 2016). Because not all of Glasser areas are involved in processing language, we restricted the analysis to 25 language-related parcels of the left hemisphere, as has been previously reported (Rolls et al., 2022). In most of the 25 areas, we found a coexistence of syntax and semantics specific voxels, the proportion of which varies in accordance to which macroscopic network the areas would most likely fall into. For instance, the posterior STG regions comprised a majority of syntax-related voxels. Conversely, regions belonging to the angular gyrus showed a greater number of semantics-related voxels. The latter result is consistent with past reports that the angular gyrus is involved in combinatorial semantics (Bemis & Pylkkanen, 2013; Price et al., 2015, 2016).

### Bridging Spatial Scales: Intracranial Evidence and 7T Resolution

Our report of a gradient of specialization per region is in line with studies using intracranial recordings, the best spatially resolved method available. Indeed, recent studies showed that electrodes distant from each other by not more than a few millimeters, can assume diverse yet critical language-related computations, whether it be for lexical and orthographic processing (Woolnough et al., 2020), phrase composition (Murphy et al., 2022), or sentence comprehension (Woolnough et al., 2023). Not only do these specialized groups of neurons respond to specific language features, but their temporal receptive fields also vary based on language processing characteristics (Regev et al., 2024).

While its spatial resolution is still lower than that of intracranial recordings, 7T fMRI also still makes it possible to go beyond the classical macroscopic functional segregation of large cortical areas in the human brain. One recent example of redescription due to higher spatial resolution concerns the Visual Word Form Area (VWFA). This brain area is part of the Ventral Occipital Temporal Cortex (VOTC) and specializes for work recognition as opposed to other types of visual stimuli. It was originally described as one continuous area of the VOTC showing increasing specialization for word recognition following a posterior to anterior gradient (Cohen et al., 2002; Vinckier et al., 2007). However, using 7T fMRI at 1.2 mm isotropic resolution, Zhan et al (2023) found that, in individual subjects, the VWFA splits into small cortical patches scattered along the gyrus, rather than a single, spatially contiguous region, with some voxels even specializing for a single written language in bilingual readers. The same type of finer-grained mapping made possible by 7T functional imaging has been applied to mathematical areas involved in calculation (Czajko et al., 2024), or visual encoding of pictures (Allen et al., 2021). By approaching the spatial resolution of classical intracranial recordings while still allowing to image the entire human brain in normal participants, ultra-high field functional MRI can provide a significant contribution to the neuroscience of language.

### Methodological Limitations

Our study presents several limitations that warrant discussion. First, while the dissociation between syntax and semantics networks is supported by multiple statistical analyses, the spatial prevalence of these networks at the individual level exhibits some variability, which may be due to a small number of trials and runs. In our first cohort (20 participants), experiment 1 rested on a single run of ∼14 minutes, and language-related Glasser areas showed activated voxels in only ∼70% of the subjects. However, in our second cohort, where the number of sentences per category was doubled, the number of activated voxels increased substantially. This observation suggests that at the high resolution afforded by 7T fMRI, a higher volume of linguistic input is necessary to reliably distinguish between syntax- and semantics-related populations at the individual level.

Second, although this study investigated the neural instantiation of syntactic movement, and showed a global increase in activation for conditions with movement versus without movement, the minimal pairwise syntactic contrasts isolating each specific movement type (wh, verb, or clitic) yielded almost no significant effects, neither at the group level nor in subject-specific voxels. While this appears as a failure to replicate previous findings (in particular that of Ben-Shachar et al., 2004; Shetreet & Friedmann, 2014), it may also be a reflection of the inherent challenges in performing group-level averaging with high-resolution 7T fMRI data. As noted by Zhan et al. (2023), the increased spatial precision of ultra-high-field imaging often reveals functional organization that is too anatomically idiosyncratic to be captured by traditional macroscopic group templates. The fact that we found significant, opposing effects of syntactic complexity—increased activation in syntax-related voxels and decreased activation in semantics-related voxels—highlights the necessity of individual-level analysis to resolve these fine-grained functional distinctions. Future studies should merely ensure that the corresponding conditions are repeated sufficient times as to achieve a stable measurement for each individual sentence.

Finally, we used semantically anomalous sentences with real words to isolate syntactic operations, rather than Jabberwocky stimuli. As explained in the Introduction, there is a strong logic in this choice, namely facilitating the syntactic parsing of those stimuli by making sure that the syntactic category and features of every word is unambiguous (which is not the case when pseudowords are used). Nevertheless, it remains possible that the use of pseudowords would yield different activation profiles. Integrating both types of stimuli in the same experimental design would be a valuable next step to further characterize the computational properties of the syntax and semantics voxels identified here.

### Conclusions

In conclusion, this 7T fMRI study provides evidence for the existence of specialized functional networks for syntax and semantics, characterized by a distinct double dissociation in their response to syntactic movement. While a sensitivity to syntactic complexity is distributed across much of the language system, our findings identify key hotspots in the posterior temporal and inferior frontal lobes that maintain a high degree of specialization for structural operations. At the individual level, the emergence of functional idiosyncrasies, identifiable only through high-field fMRI at 7T, suggests that the language network is composed of intermingled, specialized populations rather than uniform macroscopic regions. These results advocate for the broader adoption of high-resolution imaging, possibly going all the way to the record field of 11.7 T (Boulant et al., 2024), and of using hierarchically organized language localizers to robustly dissect the functional architecture that support human language processing.

## Methods

### Ethics

Participants provided written informed consent for the fMRI study and received monetary compensation. The study was approved by the local ethics committee in the NeuroSpin Center (CPP 100055), and the study was conducted in accordance with the Declaration of Helsinki.

### Participants

The experiment was advertised on the laboratory recruitment platform. Participants had to meet the following criteria in order to be considered eligible: being French native speakers; being between 18 and 45 years old (to avoid age-related vascular changes affecting the BOLD signal); not taking psychoactive drugs; not being pregnant; having normal or corrected to normal vision; right-handed; not having metal implants in the body. 20 adult participants were recruited (8 women, 12 men; mean age: 26.1 ± 7.6). All participants performed one run of the RSVP truth judgement task and four runs of the single-glance mini-sentences task.

### Replication cohort of independent participants

The replication participants were enrolled for a large-scale individual brain mapping study largely exceeding the scope of this study and formed a completely independent group from the 20 participants mentioned above. One experimental session consisted of five runs of the same RSVP truth judgement task with a larger number of conditions including both mathematical and non-mathematical statements (same protocol as in Moreno et al). We restricted the analysis to conditions matching that of the present study.

### Data acquisition

Brain images were acquired using a 7-T Magnetom scanner (Siemens, Erlangen, Germany) with a 1Tx/32Rx head coil (Nova Medical, Wilmington, USA) at the NeuroSpin Center of the French Alternative Energies and Atomic Energy Commission. Di-electric pads were used for all 20 participants). In order to keep head movements to a minimum, a tape was attached to the forehead of each participant and the head coil to provide tactile feedback to the participants whenever they attempted to move their head. To minimize light reflections inside the head coil, a piece of black paper was inserted to cover the inner surface of the transmitter coil element. Stimuli were presented on a BOLDscreen 32 LCD screen (Cambridge Research Systems, Rochester, UK; 69.84 × 39.29 cm, resolution = 1920 × 1080 pixels, refresh rate = 120 Hz, viewing distance = ∼200 cm), at the head-end of the scanner bore. Participants viewed the screen through a mirror attached to the head coil. The entire scanning session lasted approximately 90 minutes.

For each participant, we acquired one run for experiment1, lasting 13 minutes and 49 seconds (due to technical issues, two runs of task 1 were acquired for sub-01) and four runs for task 2 (lasting 11 minutes and 43 seconds each). Functional data were acquired with a two-dimensional (2D) gradient-echo echo-planar imaging (EPI) sequence [repetition time (TR) = 2000 ms, echo time (TE) = 21 ms, voxel size = 1.2 mm isotropic, multiband acceleration factor = 2; encoding direction: anterior to posterior, iPAT = 3, flip angle = 75, partial Fourier = 6/8, bandwidth = 1488 Hz per pixel, echo spacing = 0.78 ms, number of slices = 70, no gap, reference scan mode: GRE, MB LeakBlock kernel: off, fat suppression enabled]. To correct for EPI distortion, a five-volume functional run with exactly the same parameters except for opposite phase encoding direction (posterior to anterior) was acquired immediately before each task run. Participants were instructed not to move between these two runs. Manual interactive shimming of the B0 field was performed prior to functional runs for all participants. The system voltage was set at 250 V for all sessions, and the fat suppression was decreased per run to ensure that the specific absorption rate did not exceed 63% for all functional runs. To minimize artifacts and increase the SNR around the Anterior Temporal Lobe (ATL), the functional data acquisition slab was placed to exclude the eyes and the ear canal signal dropout region, so that the VOTC, especially the anterior OTS above the ear canal, was covered as much as possible.

High-resolution MP2RAGE anatomical images were obtained after 1 run of task 1 and 2 runs of task 2 (resolution = 0.65 mm isotropic, TR = 5000 ms, TE = 2.51 ms, TI1/TI2 = 900/2750 ms, flip angles = 5/3, iPAT = 2, bandwidth = 250 Hz/Px, echo spacing = 7 ms).

### Experimental design

#### Functional tasks

##### Experiment 1 (RSVP truth judgment)

20 participants performed one functional run with 100 sentences, adapted from Moreno et al. (2025), divided into 5 conditions of increasing complexity: Lists of consonant strings, lists of words, semantically anomalous sentences, generic meaningful sentences, and contextual meaningful sentences (see figure 1A for examples). The 4 categories using real words were defined as follows:

- **Word Lists:** Pseudo-randomized strings of content words (nouns, verbs) and closed-class words (prepositions, determinants). These served as a baseline to provide lexical input while preventing syntactic parsing or semantic composition.
- **Semantically Anomalous ("Meaningless") Sentences:** Syntactically correct sentences where content words were replaced by unrelated words of the same lexical category (e.g., *"The flag is an old analysis…"*). These maximized syntactic processing while minimizing sentence-level meaning. Of the 40 sentences, half of them were true (T), and half were false (F).
- **Factual Knowledge:** Meaningful sentences concerning generic, context-independent facts (e.g., *"The sun rises in the east"*). These were used to identify regions involved in basic compositional semantics and world-knowledge verification.
- **Contextual Knowledge:** These stimuli introduced a specific geographical, historical, or social frame (e.g., *“In Japan…”*) to determine the truth value of a subsequent proposition. To ensure participants integrated the context, we employed a 2×2 balanced design crossing **Contextual Truth** (True vs. False within the frame) with **Out-of-Context Plausibility** (True vs. False in general). This resulted in four equal subsets of 10 stimuli each: **T/T** (True in context/True generally, e.g., *“In Beijing, pollution is a major problem”*), **T/F** (True in context/False generally, eg. *“In volcanic islands the sand is black”*), **F/T** (False in context/True generally, eg. *“In the Amazon spiders are harmless animals”*), **F/F** (False in context/False generally, eg. *“In Kenya the lakes of the plains freeze regularly”*) This design forced a reliance on specific contextual integration rather than immediate heuristic judgments.

The participant’s task was to judge whether the semantic content of each sentence was true or false. As a convention, when the sentence had no semantic content (e.g. for lists of words), we instructed the participants to answer false. For each condition, we randomly assigned 20 sentences from a pool of 40, while ensuring that the proportion of true and false sentences was balanced when applicable. The number of words was matched between conditions (mean = 8.93). The sentences were presented in a Rapid Serial Visual Presentation fashion. Each trial started with a red alerting cross for 350ms, followed by a word-by-word presentation (350ms per word). The Mean duration of the sentences was 3200ms (+/- 350ms). The response and rest period had a mean duration of 4450ms. Stimulus Onset Asynchrony was 8000ms on average, with some jitter (SOA = 8000 + 100*Poisson(10)) such that 64% of SOAs fell between 7400 and 8600ms. No feedback was provided for this run.

##### Replication Cohort

We analyzed the first 13 participants from the iCortex longitudinal study conducted at NeuroSpin, who performed the same RSVP truth judgement task on a larger number of language and mathematical conditions (stimuli from Moreno et al., 2025). We restricted the analysis to the 5 above conditions. Participants from the replication cohort were exposed to 40 sentences per condition, i.e. twice as much as above. All other experimental parameters were identical between groups.

##### Experiment 2 (single-glance mini-sentences)

Participants performed four functional runs, grouped by pairs, with a total of 240 stimuli being presented (120 stimuli x 2 repetitions). We designed the stimuli such that, across conditions, all stimuli should : i) have exactly 3 syllables; ii) have the same average length between conditions (mean = **11.3 characters**, including spaces and punctuation); iii) have almost the same number of letters (mean = **8.3**); iv) be randomly presented in two font sizes (36 and 48), to introduce greater variability at the retinotopic input level. The design included 10 conditions, including 3 controls and 7 grammatical conditions (see figure 1B for examples). The 3 control conditions were 1) single words, 2) lists of 3 words and 3) ungrammatical sentences, made of the same words as affirmative sentences, but presented in verb-object-subject (VOS) order, the only 3-word order that can never form a grammatical sentence in French. The 7 grammatical conditions were: 4) affirmative sentences (SVO), 5) affirmative sentences with clitic movement (SOV), 6) yes-no questions without movement (SVO+question mark ?), 7) yes-no questions with verb movement (VSO?), 8) wh-in situ questions (SVO?), 9) pure wh-movement questions (OSV?), and 10) wh-and-verb-movement questions (OVS?). We designed grammatical conditions with pairwise minimal syntactic manipulations (affirmative sentences +/- clitic movement, yes-no questions +/- verb movement and wh questions +/- wh and/or verb movement) in accordance with current linguistic theories on syntactic movement, i.e., internal merge (Rizzi, 2012). The sentences were presented in a Rapid Parallel Visual Presentation fashion, meaning that the entire stimulus was presented in a single-glance for 200ms. We chose this duration to prevent eye-saccades and ensure ‘single-glance’ processing. The trials started with an alerting red cross for 350ms, followed by visual presentation of the stimulus for 200ms and a resting period of 3650ms. Stimulus Onset Asynchrony was 4200ms on average with jitter (SOA = 4200 + 159*Poisson(10) such that 85% of SOAs fell between 3600-4800ms.

Because these mini-sentences were devoid of truth value, we could not use the same task as in Experiment 1. Instead, participants performed a one-back task. On rare trials, which were modeled but not analyzed for activation (12 or 13 trials per run), two stimuli were presented, separated by a horizontal bar. One stimulus was an exact match to the one which had just been presented, while the other differed either by a single word within the same category or belonged to a different category altogether, with the exception of single words for which the confound was a randomly selected different single word. On these trials, participants had to click on the side of the stimulus matching the immediately previous trial. Visual feedback was provided after the response (word *Correct* written in green, word *Incorrect* written in red or alerting message if the response exceeded the available 1500ms response window).

##### Behavioral task on mini-sentences

Immediately following the scanning sessions, each participant performed a simple grammatical decision task on each of the 240 mini-sentences stimuli described above, with the same single-glance RPVP presentation. On each trial, participants had to decide whether the sentence was grammatically correct in French. They were told that, as a convention, sentences where only the hyphen was missing, as in “où vas tu?” (where do you go?) which should technically be spelled “où vas-tu?”, were considered grammatically correct. We report average error rate and median reaction time over 20 participants.

### Data analysis

#### Data preprocessing

Results included in this manuscript come from preprocessing performed using *fMRIPrep* 22.0.2 (Esteban et al., 2019), which is based on *Nipype* 1.8.5 (Gorgolewski et al., 2011).

#### Statistical analysis

All the analyses presented in the study were conducted using nilearn 0.10.0, a neuroimaging python package based on scikit-learn (Abraham et al., 2014).

##### fMRI GLM Models

fMRI first-level models were estimated for each subject to model task-related blood-oxygen-level dependent (BOLD) responses. For experiment 1, the design matrix included five regressors for the onsets of the five experimental conditions: lists of consonant strings, lists of words, semantically anomalous sentences, generic meaningful sentences, contextual sentences. Activation duration was set to xxx ms before convolution with the standard hemodynamic response function. For experiment 2, ten regressors modeled the ten experimental conditions: single words, lists of three words, ungrammatical sentences, affirmative sentences, clitic movement, yes-no questions, yes-no questions with verb movement, wh-in situ, pure wh-movement and wh with verb movement. Duration was set to display duration, i.e. 200 ms. One additional regressor was included to account for the key press (index or middle finger of the right hand). Regressors of no-interest included six motion corrections to account for head translation and rotation along the three axes, as well as two regressors for physiological noise: average cerebrospinal fluid signal, and average white matter signal. To account for low-frequency signal drifts, polynomial drift models from constant to 2^nd^ order were added. Finally, a constant term was also included to model the baseline signal. Single-subject analysis was performed using a smoothing applied at a full-width half-maximum (FWMHM) of 2.4mm (twice the voxel size).

Group-level analysis was performed using a second-level model with spatial smoothing applied at a full-width half-maximum (FWMHM) of 6mm. Unless indicated, all brain activation results are reported using a voxel-wise threshold of p < 0.001, corrected for multiple comparisons across the whole brain using false discovery rate (FDR) at *α* < 0.05 and a cluster-extent threshold of k≥4 voxels.

##### Projection of Glasser atlas on native T1 image of each participant

To allow for automated anatomical labelling of patches of activation, we used a custom-built python script to project the atlas defined in Glasser and Van Essen to the native anatomy of each participant (Glasser et al., 2016). The script used functions from Freesurfer (Dale et al., 1999; Fischl et al., 1999, 2002). We first mapped the fsaverage parcellation of the atlas to the anatomy of each subject (SurfaceTransform() function in Freesurfer) and then projected this individualized parcellation back to the volume space of each subject (FSCommand(command=’mri_aparc2aseg)).

##### Spatial Overlap Analysis (Dice Similarity Coefficient)

To quantify the degree of spatial segregation between syntactic and semantic networks at the individual level, we calculated the Dice Similarity Coefficient (DSC) for each participant. This analysis was specifically conducted on the iCortex replication cohort (n=13), as these individuals provided double the amount of functional data compared to the discovery cohort, ensuring the high signal-to-noise ratio necessary for precise individual-level mapping. For each subject, we generated binary activation masks for the syntactic contrast (Colorless sentences vs. Word lists) and the semantic contrast (Factual sentences vs. Colorless sentences). These masks were thresholded using a voxel-wise Z>3.29 (p<0.001, uncorrected) and a cluster-extent threshold of k≥4 voxels, matching the parameters used for individual visualizations. The DSC was defined as: 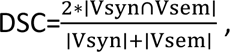 where ∣Vsyn∩Vsem∣ represents the number of voxels shared by both masks, and ∣Vsyn∣ and ∣Vsem∣ represent the total number of active voxels in the syntactic and semantic masks, respectively. The resulting coefficient ranges from 0 (no spatial overlap) to 1 (perfect spatial identity). Group-level results are reported as the mean DSC ± standard deviation across the 13 iCortex participants.

##### Regions of Interest (ROI) from task 1

We applied the functional ROI analysis method to study the activations of cortical patches isolated from task 1 contrasts. We used two main contrasts of interest: 1) Semantically anomalous sentences – lists of words (*Syntax contrast*) and 2) General sentences – Semantically anomalous sentences (*Semantics contrast*). We selected voxels responsive to these contrasts using an uncorrected voxel-wise threshold of p < 0.001 and smoothing applied at a full-width half-maximum (FWMHM) of 2.4mm. We only selected voxels that belonged to clusters of size ³ 4 (nilearn cluster_threshold parameter). To increase sensitivity, we did not use correction for multiple comparisons at this stage. Using the personalized Glasser atlases available for each subject (see section above), we restricted the subsequent analysis to Syntax and Semantics cortical patches that belonged to 25 language areas, as in Rolls et al (Rolls et al., 2022). The estimation of beta value was performed at the native resolution of our functional images (1.2mm isotropic), i.e. without any smoothing. Thus, for each participant, we had a set of beta values estimated at the native resolution for each of the 15 conditions (5 from **experiment** 1 and 10 from **experiment** 2), in 25 Glasser language regions, for 2 types of patches (Syntax patches and Semantics patches).

##### Cross-validation of Syntax and Semantic ROIs

To evaluate the functional specificity and reproducibility of the identified cortical patches, we performed a 5-fold cross-validation analysis using the iCortex replication cohort (n=13). For each participant, who performed five runs of experiment 1, we utilized a "leave-one-run-out" procedure. A first-level GLM was estimated using four of the five runs to define individual syntax and semantic ROIs. Unbiased beta values for the five experimental conditions were then extracted from the remaining, independent run. This process was repeated five times, and the resulting beta values were averaged across folds for subsequent statistical modeling.

##### Statistical framework

Statistical analysis of the extracted beta values was conducted using Linear Mixed-Effects (LME) models implemented via the *statsmodels* library in Python (v0.14+), supplemented with weighted paired *t*-tests, sign tests, and a Cochran’s *Q* heterogeneity test where indicated. LMEs were chosen for their ability to handle unbalanced designs and missing observations — occurring when specific participants lacked syntax or semantics patches in a given anatomical region — without list-wise deletion or mean imputation.

All LME and *t*-tests applied classical weighted least squares (WLS) with weights equal to the number of voxels in the patch (*n*_voxels_, linear), matching the per-region weighting used to construct Figure 6. Concretely, in each LME the response *y*, the fixed-effect design matrix *X*, and the random-effect design matrix *Z* were each premultiplied by √*n*_voxels_ prior to fitting. This is mathematically equivalent to R’s *lme4::lmer(…, weights = voxel_count)* and was verified to recover identical chi-square statistics on the headline tests (Supplementary Methods, Section S2). Models specified a random slope for voxel-type per subject (*∼ contrast_id | subject*), with automatic fallback to a random intercept when convergence failed. Wald tests on the relevant interaction terms produced χ^2^-distributed statistics; *p*-values from per-region or per-cluster tests were corrected for multiple comparisons using the Benjamini–Hochberg FDR procedure, applied independently within each test family.

##### Global and per-region voxel-type × condition analysis

To assess the functional dissociation between syntax and semantic voxels, we first fit a global LME pooling across regions, with Condition and Voxel-type as fixed factors, their interaction as the test of interest, and Region included as a fixed-effect covariate; analyses were restricted to subject × region cells containing both voxel types. The same model was then fit separately within each of the 25 testable Glasser language regions in iCortex and the 24 testable regions in Main (TGv_L was dropped from iCortex and TGv_L/STGa_L from Main, for lack of any voxel of one type) and within each of eight functionally defined clusters (with *region_name* included as a fixed-effect covariate in the cluster-level fits). For each region and cluster we additionally reported *n*_dual_, the number of participants contributing at least one voxel of each type, which gauges the extent to which the interaction is supported by within-subject (rather than between-subject) variability.

Heterogeneity of the voxel-type × condition interaction across regions — the interpretable form of a Region × Voxel-type × Condition triple interaction on unbalanced data — was assessed using Cochran’s *Q* on the per-region two-sided *z*-transformed *p*-values, weighted by *n*_dual_. The same global and per-region LMEs were additionally re-fit on the grammatical subset of the Main dataset (conditions 04–10) to test for an effect of syntactic structure within well-formed sentences.

##### Region-wise comparison of the two critical conditions in iCortex

To characterise the contribution of the semantically anomalous condition to the iCortex dissociation, we performed a systematic region-wise comparison between the two critical hierarchical conditions of Experiment 1: semantically anomalous sentences (condition 3) and generic meaningful sentences (condition 4). For each Glasser region and separately for syntax-tuned and semantic-tuned voxel sets, we computed a paired weighted *t*-test on the within-subject 3 − 4 difference, with weights equal to each subject’s *n*_voxels_ in that region (normalised to mean 1) and *df* = *n* − 1. The weighted statistic was *t* = *m̄*_w_ / √(*s*^2^_w_ / *n*), where *m̄*_w_ = Σ*w*_i_ *d*_i_ / Σ*w*_i_ and *s*^2^_w_ = Σ*w*_i_(*d*_i_ − *m̄*_w_)^2^ / Σ*w*_i_. FDR correction was applied across the 25 regions independently for each voxel-type family.

##### Sensitivity to syntactic movement

The analysis of sensitivity to syntactic movement was carried out at three levels.

###### Region-level paired contrasts in three key ROIs

In L_44, L_45, and L_STSvp, we computed weighted paired *t*-tests on syntax voxels (using the same weighted statistic as above) for four contrasts targeting individual movement types: clitic movement against basic affirmatives (05 − 04), yes-no questions with verb movement against those without (07 − 06), pure wh-movement against wh-in-situ (09 − 08), and the addition of verb movement to wh-movement (10 − 09).

Each contrast was FDR-corrected across the 3 regions in which it was computed. Supplementary Table S3 reports the *p*-values for all 25 Glasser regions.

###### Sign test on six baseline contrasts

For the broader syntax network, we re-parameterised each grammatical condition (05–10) as a within-subject difference against the simplest baseline (condition 04) computed on syntax voxels. Because the six contrasts share a common reference and are therefore correlated, we combined them via a distribution-free sign test on the number of positive contrasts per region and per cluster, with a one-tailed null probability of 0.5 per contrast (so that obtaining 6/6 positives carries an exact *p* = 1/64 ≈ .016). Sign-test *p*-values were FDR-corrected across regions and across clusters.

###### Binary movement × voxel-type LME

We coded each grammatical condition as carrying either no syntactic movement (conditions 04, 06, 08) or at least one movement (conditions 05, 07, 09, 10), and fit an LME with Movement (0/1) and Voxel-type as fixed factors, their interaction as the test of interest, Region as a fixed-effect covariate, a random slope for voxel-type per subject, and classical WLS weights = *n*_voxels_. The model was fit pooled across regions and then re-fit independently in each of the 22 testable Main regions and each of the eight functional clusters. From each fit we extracted the interaction χ^2^(1), the *semantic slope* (the coefficient of Movement at the reference level of Voxel-type) and the *syntax slope* (semantic slope + interaction, with its standard error computed from the joint covariance of the two coefficients). *p*-values for the interaction and for each of the two slopes were FDR-corrected across regions and across clusters within their respective families.

## Acknowledgment

This research was supported by INSERM, CEA, Collège de France, Sorbonne Université, ERC grant “NeuroSyntax” to S.D., and the *Fondation pour la Recherche Médicale* foundation, FDM program, grant number FDM202206015336 to T.D.B. We are grateful to all support cells at NeuroSpin for their help with subject recruitment, scanning and data processing, and to the CMRR lab in Minneapolis for their multiband EPI sequence.

## Supplementary Materials

### Divergent Neural Profiles for Mini-Sentences

We evaluated group-level brain responses to the single-glance mini-sentences using two primary contrasts (Fig. S9). The first contrast, comparing the simplest declarative sentences (Condition 4) to word lists (Condition 2), revealed two distinct response profiles within the language network. First, we observed significant signal reductions (deactivations) in syntax-sensitive regions identified in the RSVP task, including the MFG (85% prevalence), IFGtri (65%), IFGorb (40%), and pSTG (85%). Conversely, increased BOLD signal was observed in semantic-related areas, specifically the AG (60%) and aSTG (35%).

A second contrast was performed between the same declarative sentences and ungrammatical strings (the "Grammaticality" contrast). The spatial pattern of signal reductions in syntax-sensitive regions was qualitatively similar to the previous contrast (MFG: 45%; IFGtri: 35%; pSTG: 60%). Crucially, however, this contrast yielded no regions with significantly increased activation. This lack of positive activation suggests that while the transition from unstructured lists to grammatical sentences engages semantic-sensitive patches, the violation of syntax in ungrammatical strings primarily affects the degree of engagement within syntax-sensitive regions rather than recruiting additional cortical resources.

### Sensitivity of regional LME results to observation weighting

The primary analysis (presented in Table 2) used √n_voxels precision weighting. Tables S1 and S1b reproduce the nested cluster/region structure of Table 2 under two alternative weightings: unweighted and linear *n_voxels*. The same overall pattern of regional and cluster-level significance was observed under all three weightings, with minor re-ordering of marginally significant regions.

**Table S1.** Voxel-type × condition interactions by cluster and region (unweighted, sensitivity analysis)

| Region | *iCortex* |  |  | *Main* |  |  |
| --- | --- | --- | --- | --- | --- | --- |
| | $\chi^2$ | *p* (FDR) | *n* dual | $\chi^2$ | *p* (FDR) | *n* dual |
| *44_L* | 13.91 | .016* | 11 | 9.68 | .533 | 6 |
| *45_L* | 10.06 | .053<sup>†</sup> | 10 | 9.59 | .533 | 7 |
| *47l_L* | 12.55 | .024* | 10 | 6.52 | .824 | 7 |
| *47s_L* | 4.45 | .363 | 7 | 19.60 | .082<sup>†</sup> | 4 |
| *STSdp_L* | 22.48 | .001** | 6 | 18.25 | .111 | 3 |
| *STSvp_L* | 28.70 | < .001*** | 9 | 25.24 | .016* | 9 |
| *STSda_L* | 15.84 | .012* | 7 | 10.98 | .533 | 3 |
| *STSva_L* | 11.60 | .034* | 8 | 10.36 | .533 | 8 |
| *TGd_L* | 4.11 | .392 | 6 | 2.10 | .990 | 1 |
| *TGv_L* | — | — | 0 | — | — | 0 |
| *STGa_L* | 9.90 | .053<sup>†</sup> | 2 | — | — | 0 |
| *55b_L* | 25.41 | < .001*** | 9 | 30.50 | .004** | 5 |
| *SFL_L* | 18.32 | .004** | 11 | 17.11 | .141 | 11 |
| *8Av_L* | 19.38 | .003** | 11 | 10.23 | .533 | 14 |
| *PGi_L* | 32.15 | < .001*** | 13 | 26.23 | .015* | 13 |
| *TPOJ1_L* | 15.43 | .012* | 5 | 22.46 | .036* | 1 |
| *PSL_L* | 14.93 | .012* | 9 | 13.07 | .348 | 4 |
| *9a_L* | 5.09 | .303 | 8 | 14.34 | .265 | 10 |
| *9p_L* | 14.00 | .016* | 10 | 2.30 | .990 | 10 |
| *8BL_L* | 13.36 | .019* | 11 | 5.68 | .841 | 9 |
| *A5_L* | 9.92 | .053<sup>†</sup> | 6 | 43.44 | < .001*** | 2 |
| *9m_L* | 15.16 | .012* | 10 | 9.41 | .533 | 15 |
| *STV_L* | 11.41 | .035* | 8 | 8.67 | .591 | 3 |
| *TE1a_L* | 8.15 | .103 | 10 | 9.88 | .533 | 12 |
| *10v_L* | 5.27 | .296 | 10 | 16.41 | .157 | 3 |
| *10pp_L* | 10.33 | .052<sup>†</sup> | 2 | 5.74 | .841 | 1 |

*Note.* Same structure as Table 2 but without observation weighting. Cluster-level rows (bold) contain region\_name as a fixed-effect covariate; constituent regions are shown below each cluster (italic, indented). Regions not assigned to any cluster are at the bottom.
<sup>†</sup> $p < .10$ . \* $p < .05$ . \*\* $p < .01$ . \*\*\* $p < .001$ .

**Table S1b.**
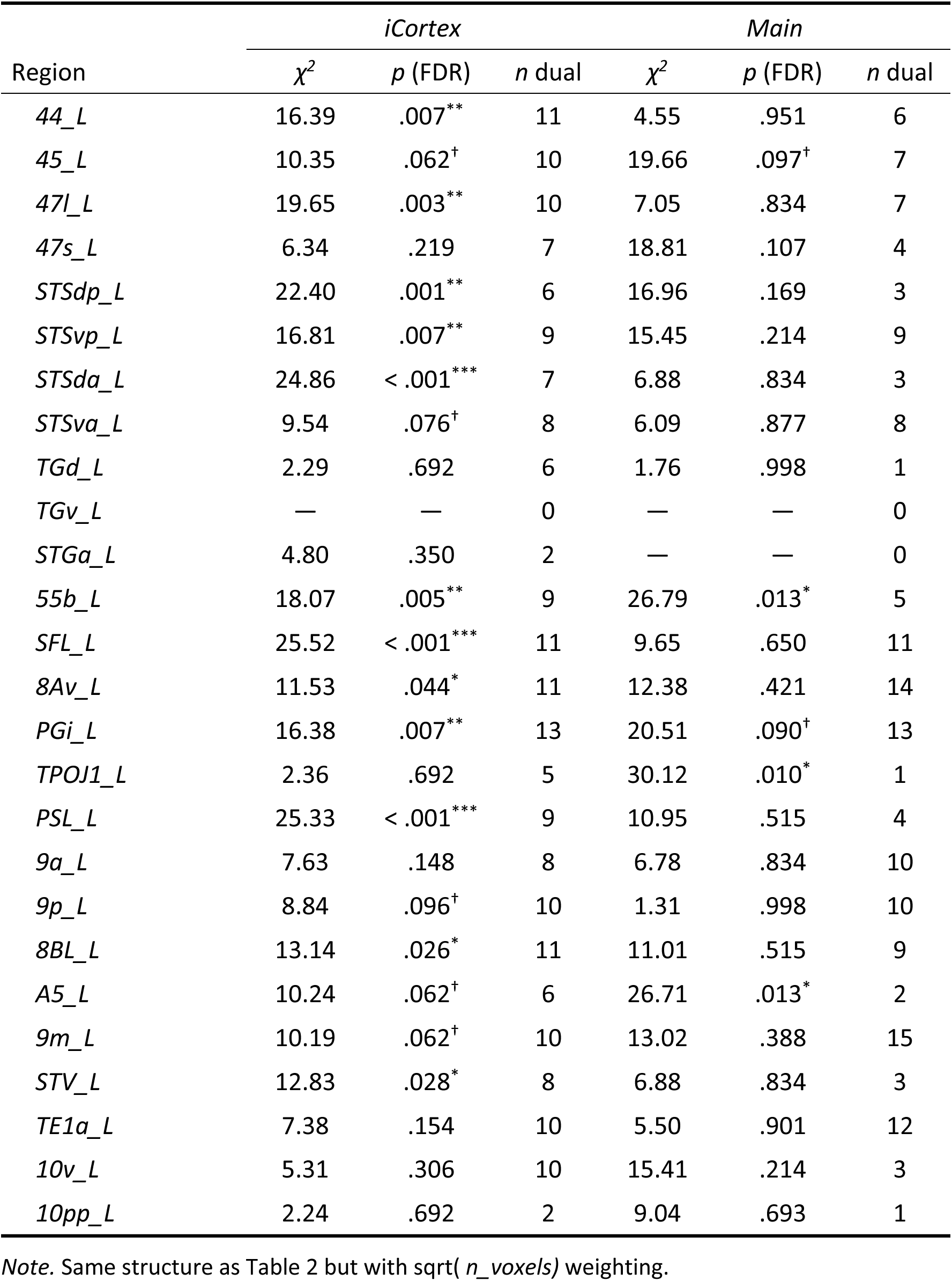
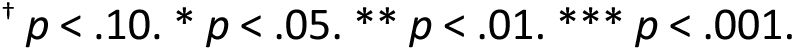
Voxel-type × condition interactions by cluster and region (sqrt(n_voxels) weighting, sensitivity analysis)

| Region | *iCortex* |  |  | *Main* |  |  |
| --- | --- | --- | --- | --- | --- | --- |
| | $\chi^2$ | *p* (FDR) | *n* dual | $\chi^2$ | *p* (FDR) | *n* dual |
| 44_L | 16.39 | .007** | 11 | 4.55 | .951 | 6 |
| 45_L | 10.35 | .062<sup>†</sup> | 10 | 19.66 | .097<sup>†</sup> | 7 |
| 47l_L | 19.65 | .003** | 10 | 7.05 | .834 | 7 |
| 47s_L | 6.34 | .219 | 7 | 18.81 | .107 | 4 |
| STSdp_L | 22.40 | .001** | 6 | 16.96 | .169 | 3 |
| STSvp_L | 16.81 | .007** | 9 | 15.45 | .214 | 9 |
| STSda_L | 24.86 | < .001*** | 7 | 6.88 | .834 | 3 |
| STSva_L | 9.54 | .076<sup>†</sup> | 8 | 6.09 | .877 | 8 |
| TGd_L | 2.29 | .692 | 6 | 1.76 | .998 | 1 |
| TGv_L | — | — | 0 | — | — | 0 |
| STGa_L | 4.80 | .350 | 2 | — | — | 0 |
| 55b_L | 18.07 | .005** | 9 | 26.79 | .013* | 5 |
| SFL_L | 25.52 | < .001*** | 11 | 9.65 | .650 | 11 |
| 8Av_L | 11.53 | .044* | 11 | 12.38 | .421 | 14 |
| PGi_L | 16.38 | .007** | 13 | 20.51 | .090<sup>†</sup> | 13 |
| TPOJ1_L | 2.36 | .692 | 5 | 30.12 | .010* | 1 |
| PSL_L | 25.33 | < .001*** | 9 | 10.95 | .515 | 4 |
| 9a_L | 7.63 | .148 | 8 | 6.78 | .834 | 10 |
| 9p_L | 8.84 | .096<sup>†</sup> | 10 | 1.31 | .998 | 10 |
| 8BL_L | 13.14 | .026* | 11 | 11.01 | .515 | 9 |
| A5_L | 10.24 | .062<sup>†</sup> | 6 | 26.71 | .013* | 2 |
| 9m_L | 10.19 | .062<sup>†</sup> | 10 | 13.02 | .388 | 15 |
| STV_L | 12.83 | .028* | 8 | 6.88 | .834 | 3 |
| TE1a_L | 7.38 | .154 | 10 | 5.50 | .901 | 12 |
| 10v_L | 5.31 | .306 | 10 | 15.41 | .214 | 3 |
| 10pp_L | 2.24 | .692 | 2 | 9.04 | .693 | 1 |

*Note.* Same structure as Table 2 but with sqrt(*n\_voxels*) weighting.
<sup>†</sup> $p < .10$ . \* $p < .05$ . \*\* $p < .01$ . \*\*\* $p < .001$ .

### Matched paired contrasts on syntax voxels (Main)

We tested each of the four classical movement contrasts on syntax voxels using weighted paired *t*-tests with the same linear voxel-count weighting used in the figures (Kish’s effective sample size for degrees of freedom). The four contrasts were: clitic movement (05 − 04), verb movement in yes-no questions (07 − 06), pure wh movement (09 − 08), and added verb movement on top of wh (10 − 09). Table S3 reports the *t* and two-sided *p*-values for each contrast in each region; no contrast survived FDR correction within or across regions, with only 55b_L approaching significance on the verb-movement contrast (*t* = 4.32, *p* = .003).

**Table S3.** Matched paired contrasts on syntax voxels, by region (Main)

| Region | *Clitic (05-04)* |  | *Verb y/n (07-06)* |  | *Wh pure (09-08)* |  | *Wh+v vs Wh (10-09)* |  |
| --- | --- | --- | --- | --- | --- | --- | --- | --- |
| | $t$ | $p$ | $t$ | $p$ | $t$ | $p$ | $t$ | $p$ |
| 44_L | 0.93 | .377 | 2.31 | .048* | 0.69 | .510 | -0.86 | .416 |
| 45_L | 0.96 | .355 | 1.15 | .274 | 0.70 | .499 | -0.67 | .514 |
| 47l_L | 0.30 | .774 | 1.53 | .160 | -0.38 | .713 | 0.48 | .644 |
| 47s_L | 0.50 | .650 | 1.37 | .253 | -0.42 | .698 | -0.27 | .805 |
| 8Av_L | 0.05 | .963 | 0.60 | .568 | 0.58 | .578 | -1.08 | .314 |
| 55b_L | 0.57 | .587 | 4.32 | .003** | 0.47 | .649 | 0.10 | .924 |
| SFL_L | 1.76 | .113 | -0.61 | .559 | 1.49 | .170 | -1.60 | .145 |
| STGa_L | 1.05 | .320 | -0.43 | .676 | 1.26 | .240 | -1.01 | .338 |
| STSda_L | 2.14 | .053<sup>†</sup> | 0.10 | .920 | 0.98 | .344 | -1.11 | .288 |
| STSdp_L | 0.37 | .717 | 2.05 | .070<sup>†</sup> | -0.47 | .651 | 0.05 | .959 |
| STSva_L | 2.10 | .062<sup>†</sup> | -1.27 | .233 | 1.09 | .301 | -0.94 | .372 |
| STSvp_L | 1.82 | .106 | 1.03 | .333 | 0.73 | .484 | -0.70 | .505 |
| PGi_L | 1.41 | .211 | -2.38 | .057<sup>†</sup> | 0.28 | .793 | -1.26 | .257 |
| TPOJ1_L | 1.16 | .294 | 2.14 | .081<sup>†</sup> | 0.60 | .574 | -0.06 | .956 |
|  | *t* | *p* | *t* | *p* | *t* | *p* | *t* | *p* |
| *A5_L* | 0.02 | .988 | 1.90 | .090<sup>†</sup> | 0.67 | .522 | -0.40 | .697 |
| *8BL_L* | -0.43 | .680 | 1.04 | .342 | 0.67 | .528 | -1.02 | .350 |
| *9m_L* | 0.08 | .937 | -0.07 | .949 | -0.09 | .929 | -0.37 | .717 |
| *9p_L* | 0.17 | .868 | -0.41 | .697 | -0.26 | .805 | -0.02 | .982 |
| *PSL_L* | 1.31 | .234 | 1.94 | .098<sup>†</sup> | 1.13 | .298 | -1.19 | .277 |
| *STV_L* | 0.36 | .727 | 1.18 | .275 | 0.05 | .960 | 0.06 | .956 |
| *TE1a_L* | 1.06 | .325 | -1.65 | .145 | -0.28 | .791 | -0.66 | .534 |
| *9a_L* | -1.08 | .322 | -0.29 | .781 | -1.81 | .122 | 0.46 | .661 |
| *10v_L* | 1.81 | .245 | -1.75 | .255 | -0.34 | .772 | -0.45 | .704 |
| *10pp_L* | -0.17 | .888 | -0.85 | .532 | -0.42 | .739 | 1.13 | .439 |
| *TGd_L* | 0.01 | .995 | -0.04 | .973 | 0.03 | .981 | -0.13 | .919 |
| *TGv_L* | — | — | — | — | — | — | — | — |

*Note.* Weighted paired *t*-tests on syntax voxels, using linear voxel-count weighting. Four contrasts test increments in movement type: clitic movement (05–04), verb movement in yes-no questions (07–06), pure wh movement (09–08), and wh + verb on top of wh (10–09). *p*-values are two-sided, uncorrected; no cell survived FDR correction across the 25 regions within any single contrast.
<sup>†</sup> $p < .10$ . \* $p < .05$ . \*\* $p < .01$ . \*\*\* $p < .001$ .

---

**Figure S1 (related to figure 2).**
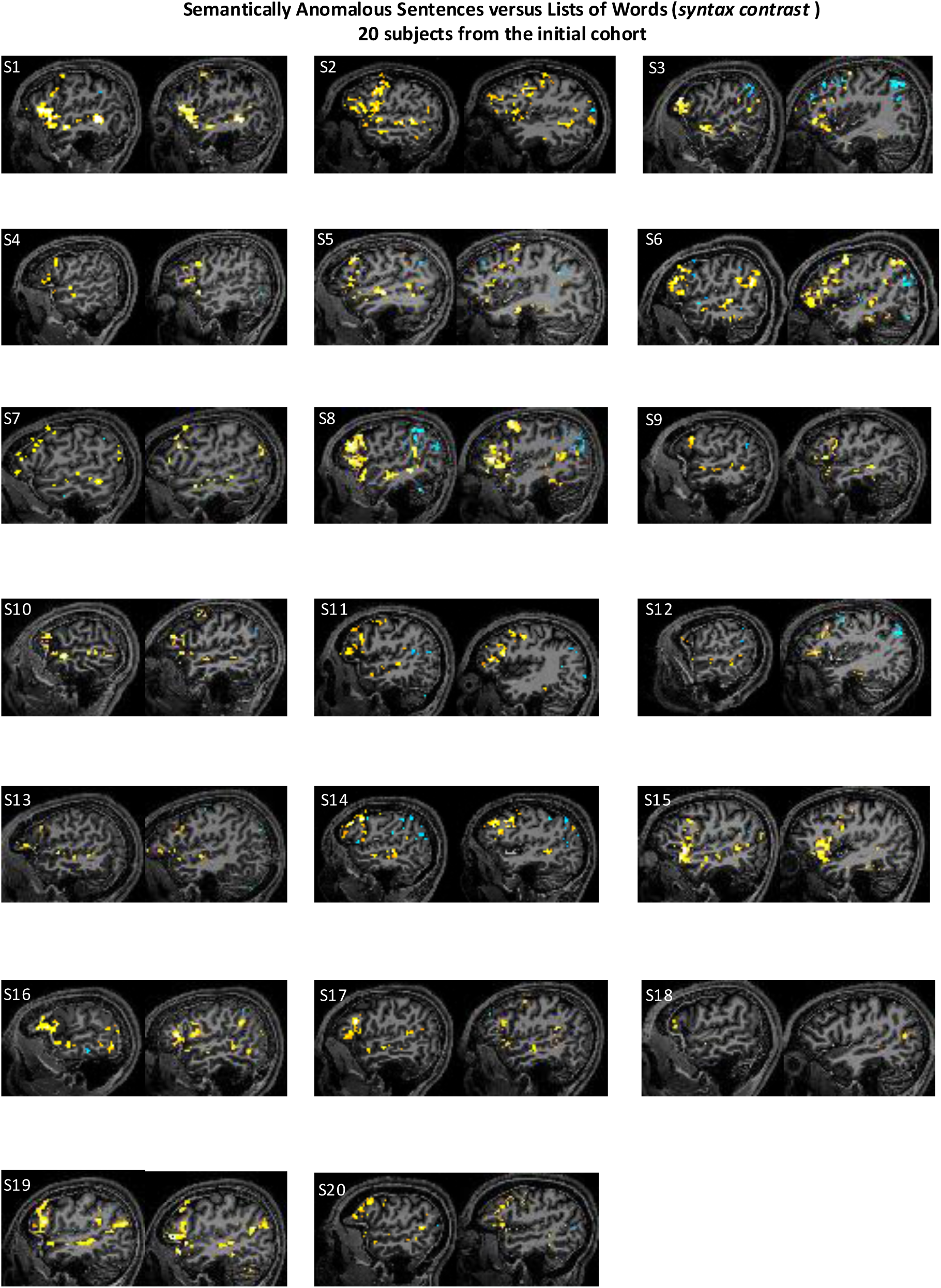
Contrast between semantically anomalous sentences and lists of words (Syntax contrast). Individual results for each of the 20 subjects of the primary cohort that performed both tasks. Uncorrected contrast maps (p<0.001 uncorrected, cluster-extent threshold of k≥4 voxels).

**Figure S2 (related to figure 2).**
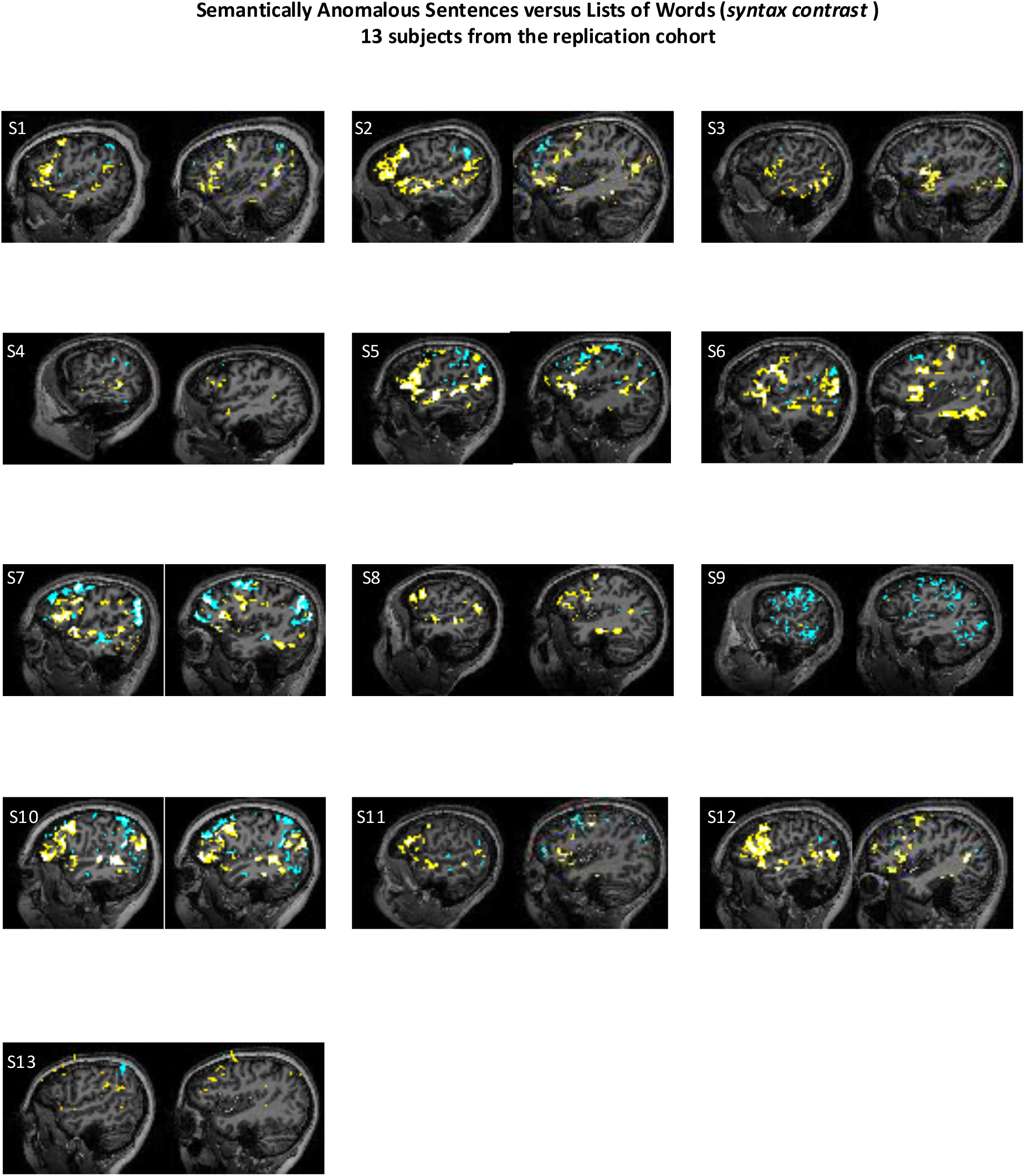
Contrast between semantically anomalous sentences and lists of words (Syntax contrast). Individual results for each of the 13 subjects from the replication cohort. Uncorrected contrast maps (p<0.001 uncorrected, cluster-extent threshold of k≥4 voxels).

**Figure S3 (related to figure 2).**
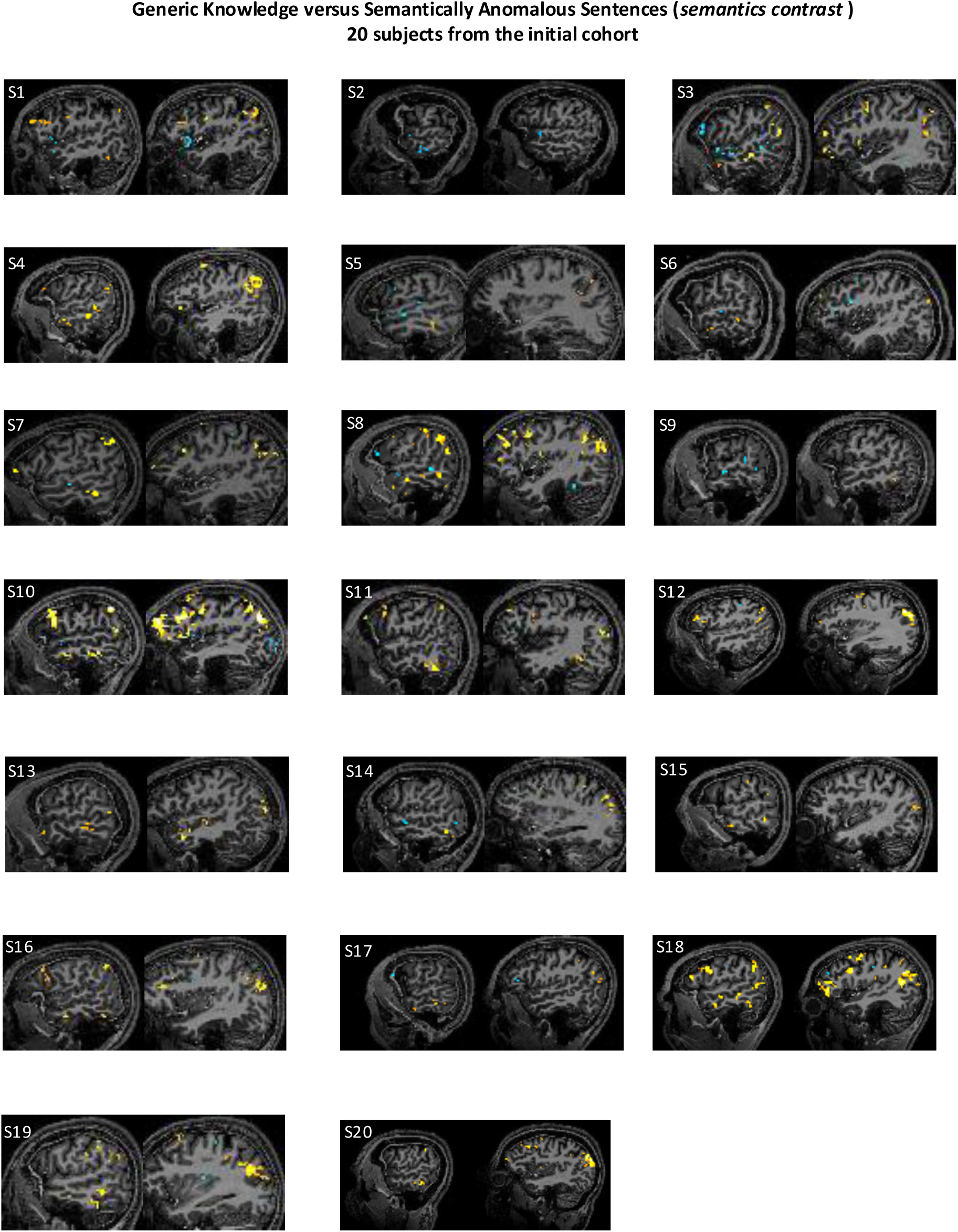
Contrast between Generic Knowledge and semantically anomalous sentences (Semantics contrast). Individual results for each of the 20 subjects of the primary cohort that performed both tasks. Uncorrected contrast maps (p<0.001 uncorrected, cluster-extent threshold of k≥4 voxels).

**Figure S4 (related to figure 2).**
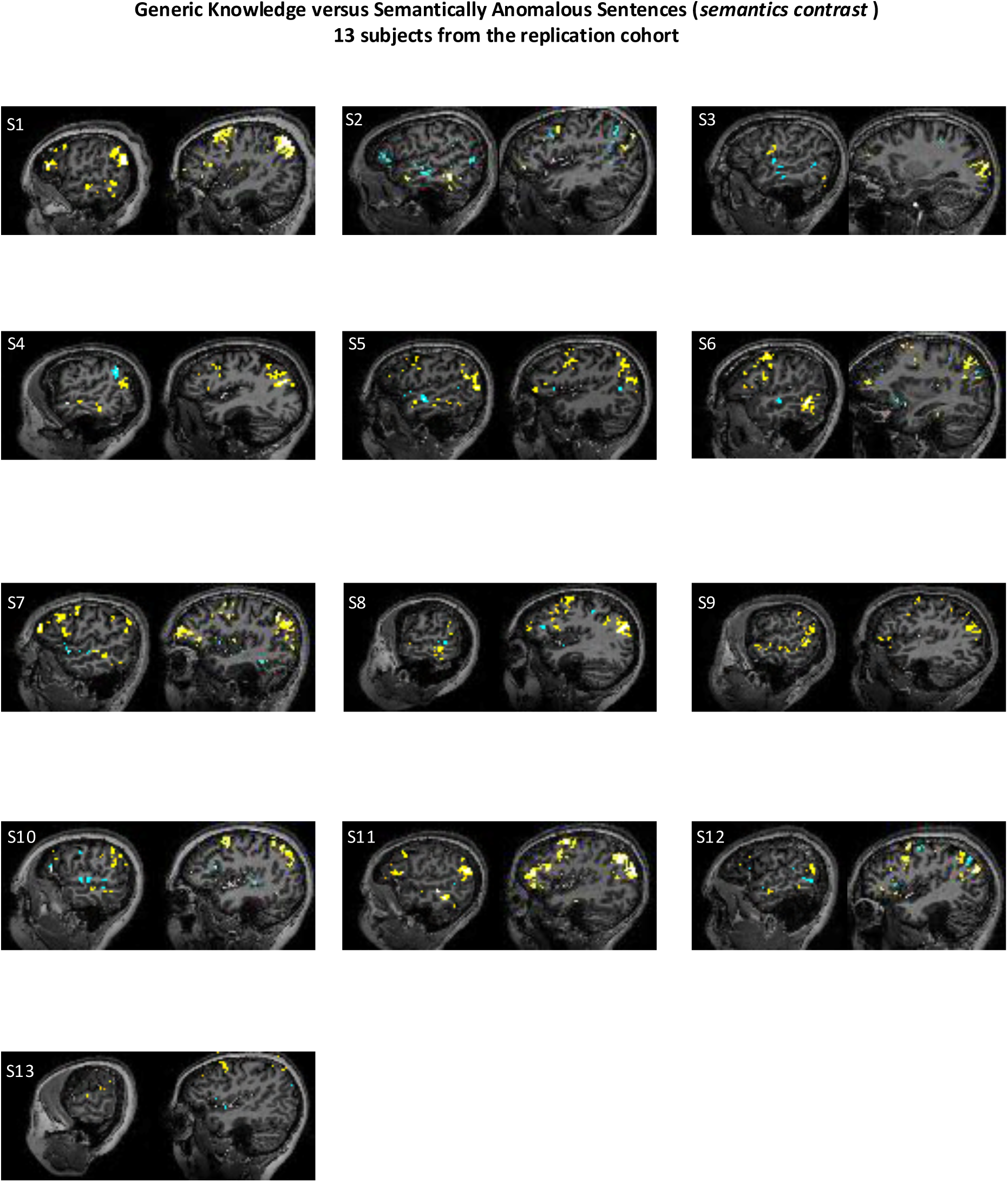
Contrast between Generic Knowledge and semantically anomalous sentences (Semantics contrast). Individual results for each of the 13 subjects from the replication cohort. Uncorrected contrast maps (p<0.001 uncorrected, cluster-extent threshold of k≥4 voxels).

**Figure S5 (related to figure 3).**
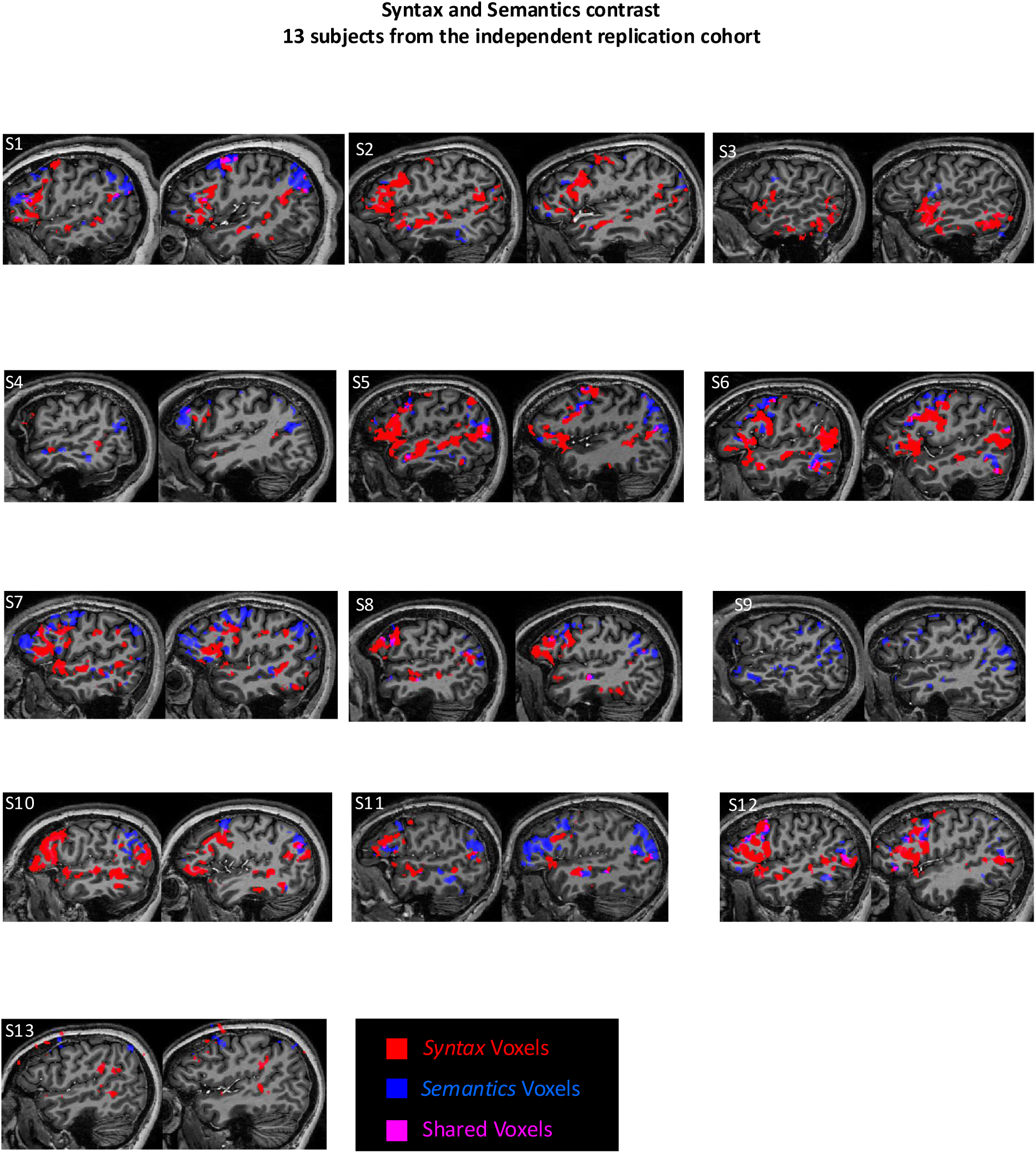
Syntax and Semantics responsive voxels superimposed on two sagittal slices of the left hemisphere, using the native T1-anatomy of all 13 participants of the replication cohort. Contrast maps presented are uncorrected, with a p-value <0.001, cluster-extent threshold of k≥4 voxels.

**Figure S6 (related to figure 5).**
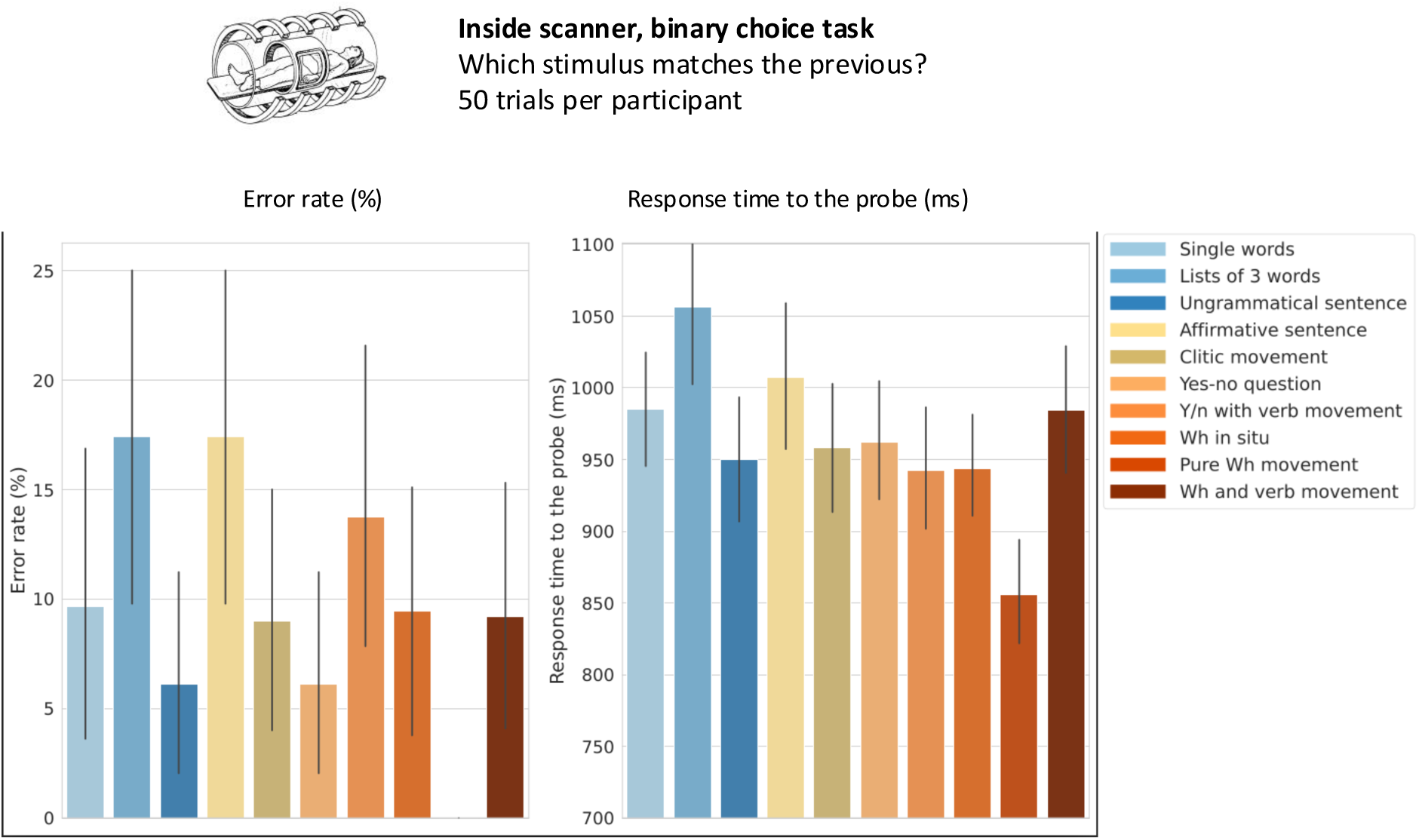
Behavioral analysis of the match-to-sample task with mini-sentences shows no effect of syntactic complexity.

**Figure S7 (related to figure 4).**
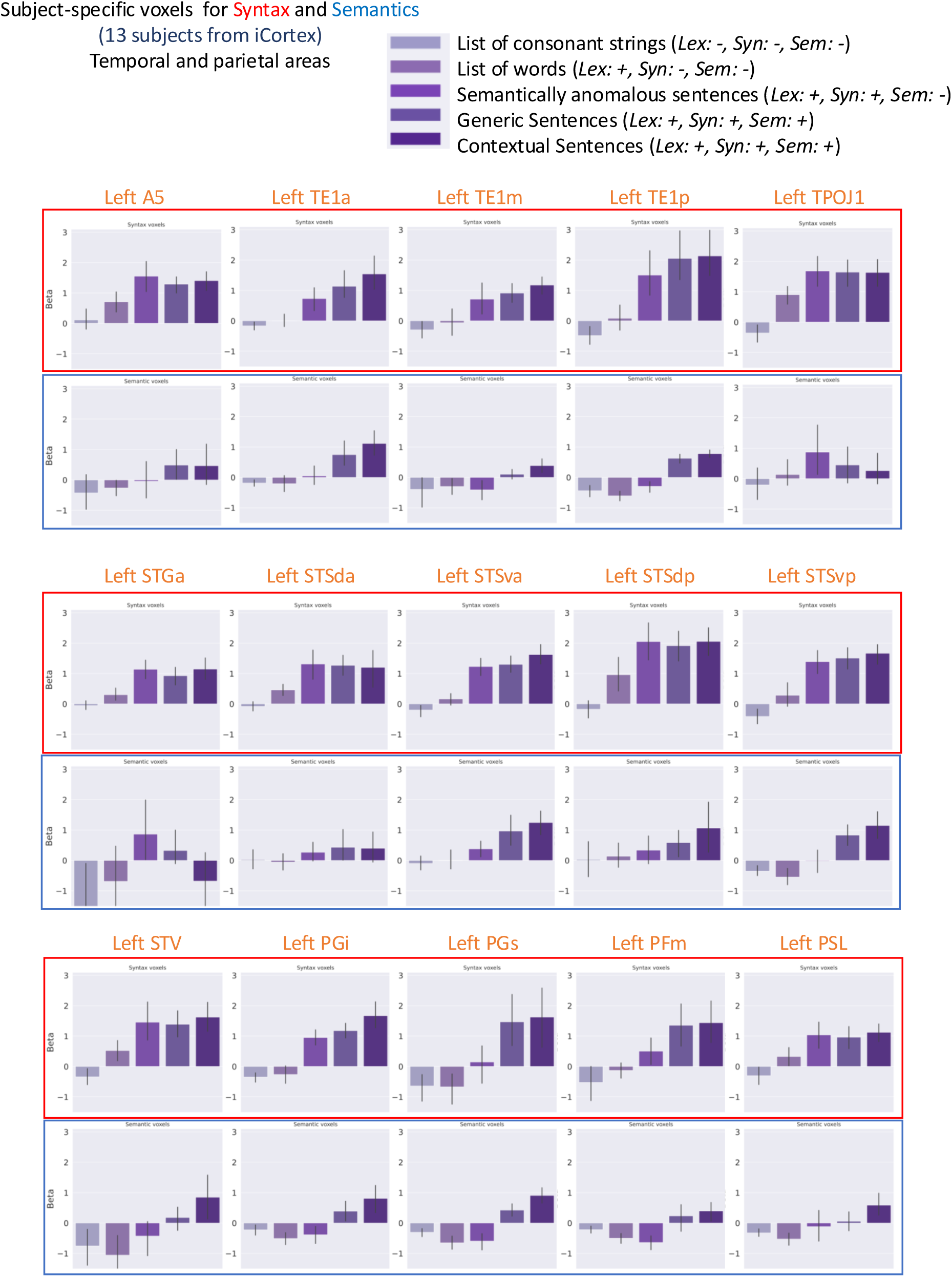
Cross-validation of the profiles of activation of Syntax and Semantics voxels. Cross-validated activation profiles of Syntax and Semantics voxels to the 5 conditions from Task 1 (RSVP truth judgement task). Temporal and parietal Glasser language regions are presented.

**Figure S8 (related to figure 4).**
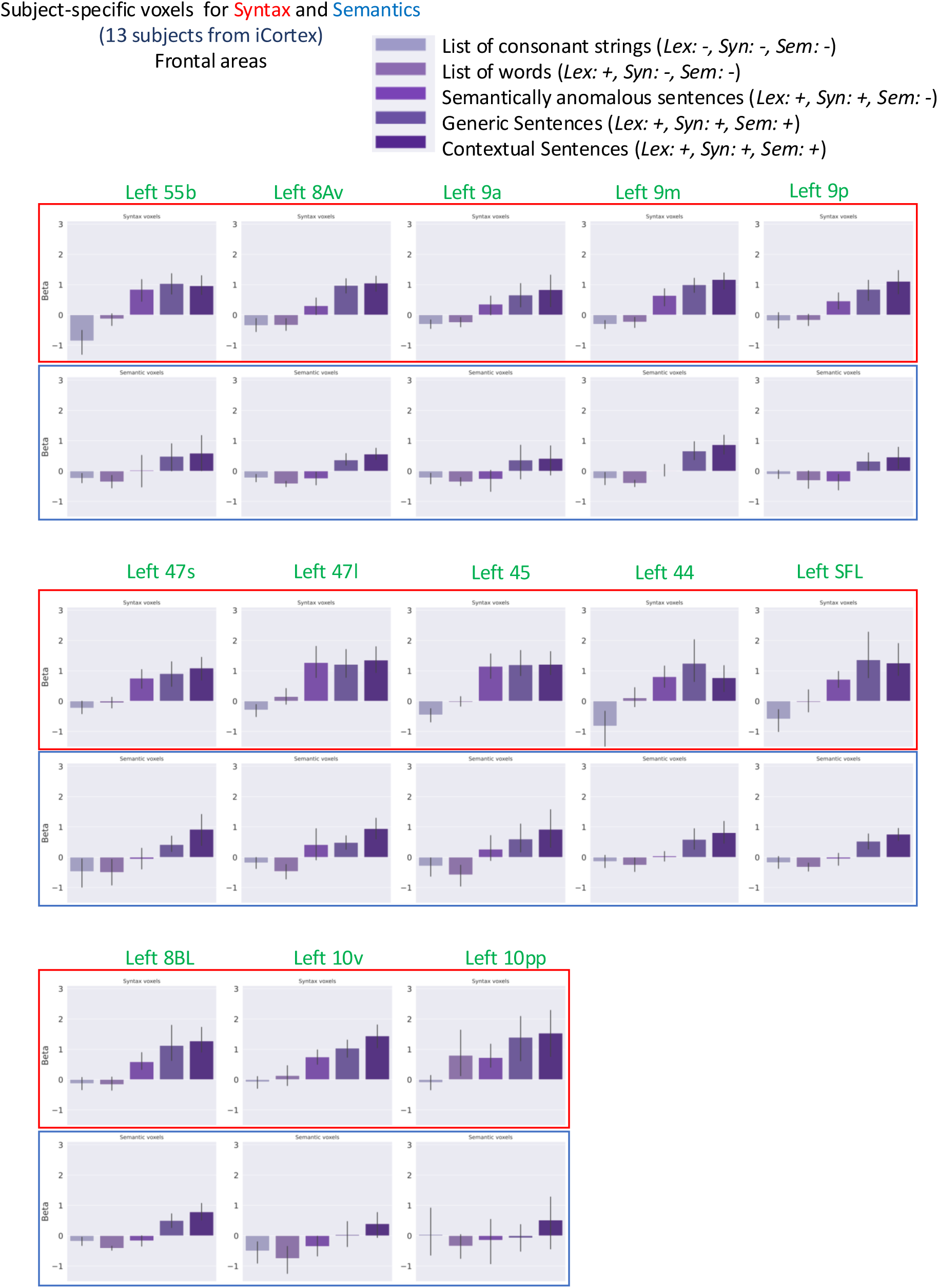
Cross-validation of the profiles of activation of Syntax and Semantics voxels. Cross-validated activation profiles of Syntax and Semantics voxels to the 5 conditions from Task 1 (RSVP truth judgement task). Frontal Glasser language regions are presented.

**Figure S9:**
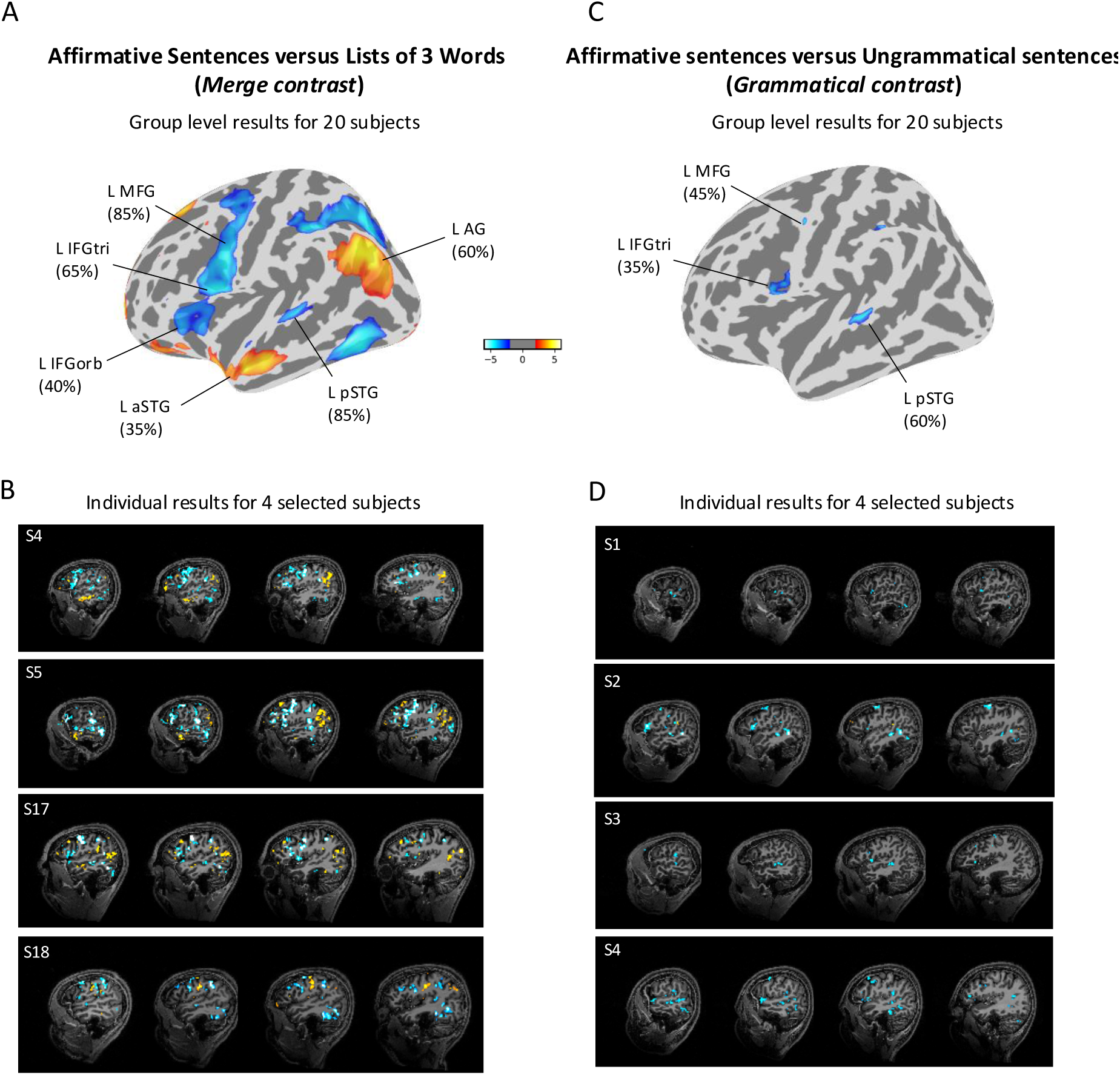
Divergent neural responses to minimal syntactic structure at group and individual levels. A: Group-level segregation of the minimal sentence core. The contrast between minimal declarative sentences (3-word affirmative) and unstructured 3-word lists identifies regions sensitive to the onset of basic syntactic and semantic processing. Results are thresholded at p<0.05, voxel-wise False Discovery Rate (FDR) corrected. B: Individual-level granularity of minimal structure processing. Representative activation maps from four individual subjects (thresholded at p<0.001, uncorrected, cluster-extent threshold of k≥4 voxels) illustrate the consistency of the functional divergence across participants. While the syntactic core (e.g., IFG and pSTG) often exhibits relative signal reductions, specialized patches in the semantic stream (AG and aSTG) show positive BOLD recruitment for the same declarative structures. C: Group-level isolation of the grammaticality effect. The contrast between declarative sentences and ungrammatical strings (3-word sequences violating French syntax) isolates the neural cost of syntactic violations. This "grammaticality" contrast consistently yields signal modulations in the syntactic core (MFG, IFG, and pSTG) without recruiting additional semantic resources, suggesting a selective sensitivity to formal structural constraints. Results are thresholded at p<0.05, voxel-wise False Discovery Rate (FDR) corrected. D: Spatial stability of individual grammaticality responses. Individual maps (thresholded at p<0.001, uncorrected, cluster-extent threshold of k≥4 voxels) for the grammaticality contrast confirm that syntax-sensitive regions are modulated by structural integrity at the single-subject level. The lack of corresponding activation in semantic-related patches (AG and aSTG) highlights a selective functional independence between formal syntactic structure building and propositional integration.

**Figure S10 (related to figure 5).**
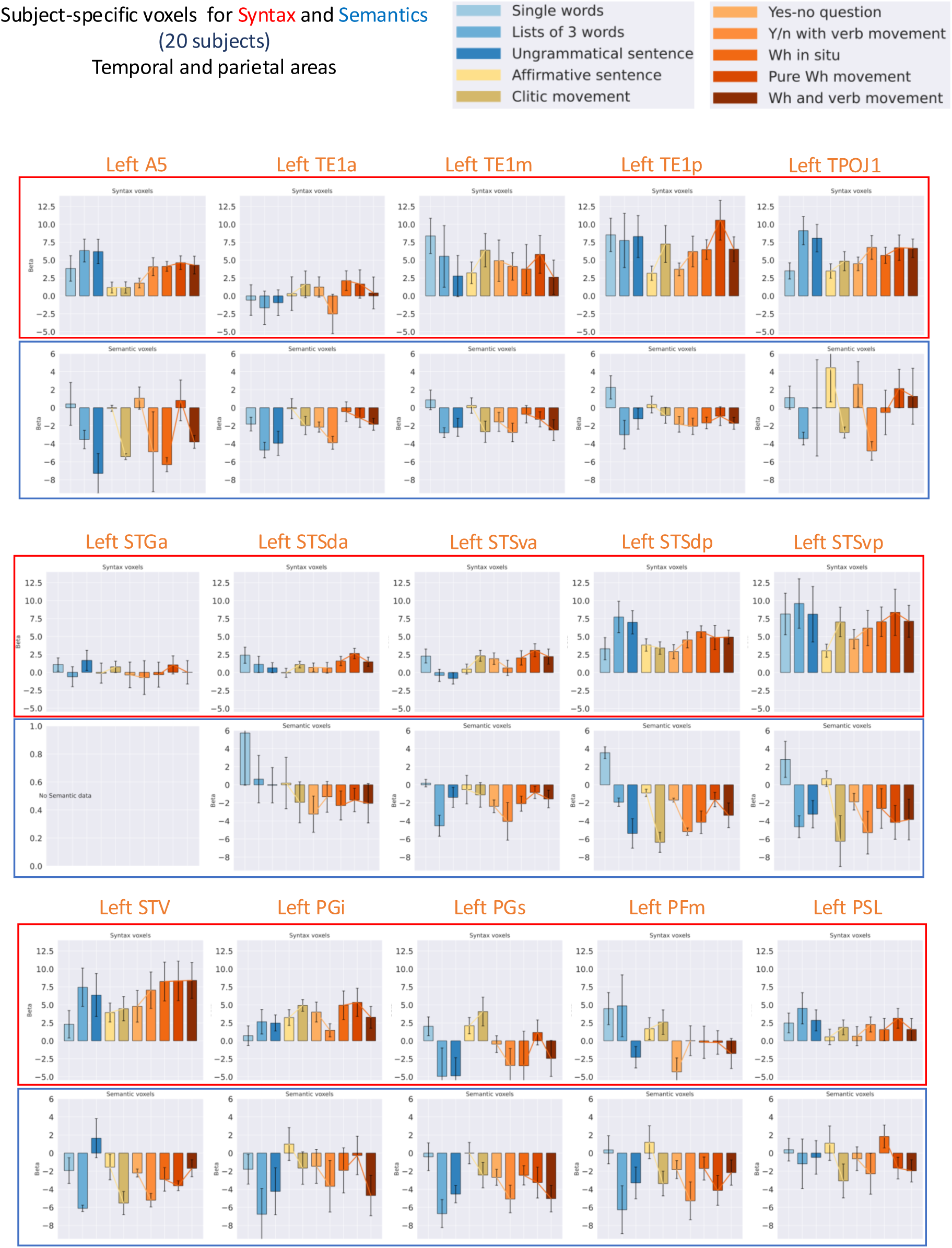
Activation profiles of Syntax and Semantics voxels to mini-sentences. Activation profiles of Syntax and Semantics voxels to the 10 mini-sentences conditions from Task 2. Temporal and parietal Glasser language regions are presented.

**Figure S11 (related to figure 5).**
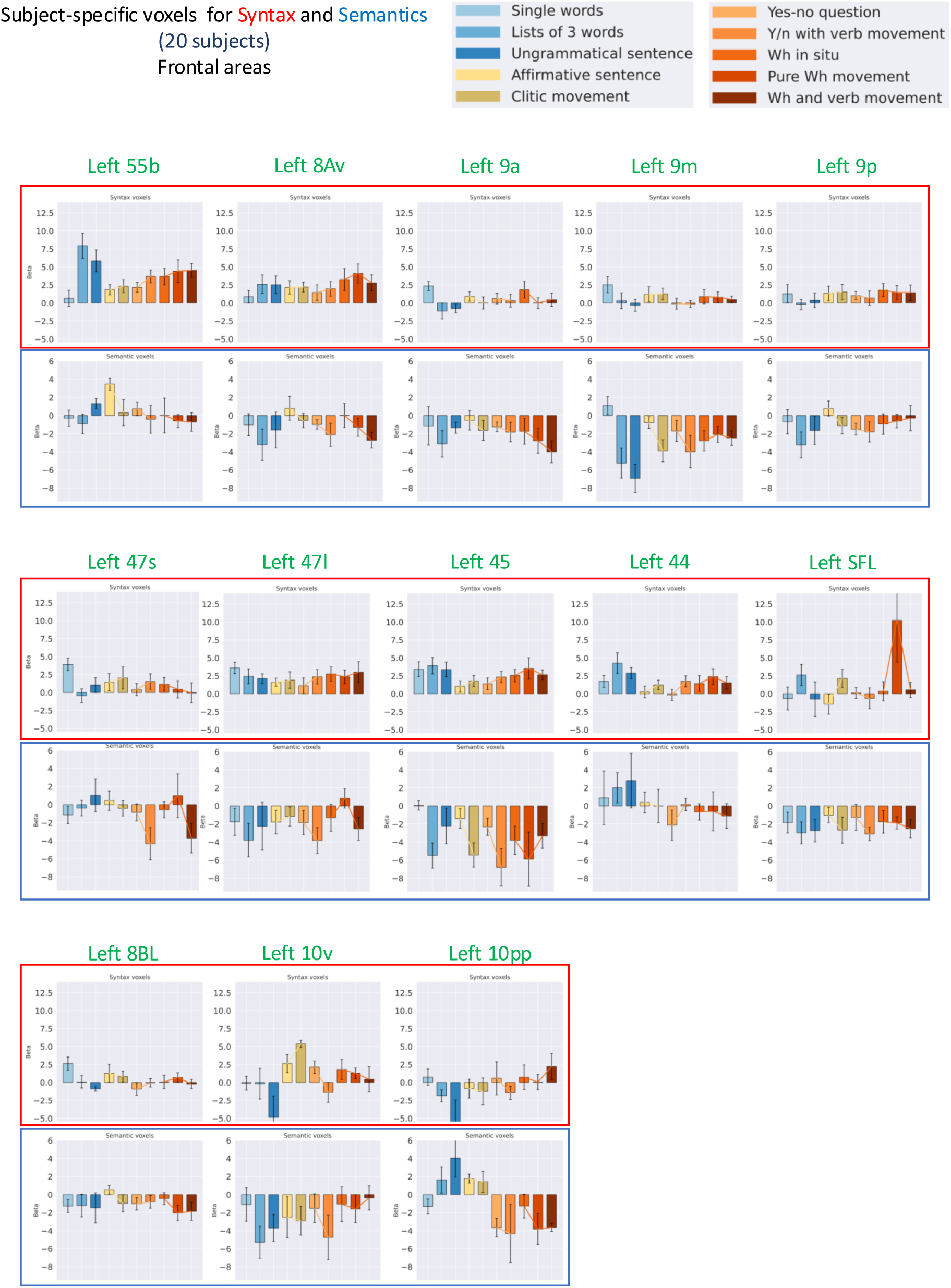
Activation profiles of Syntax and Semantics voxels to mini-sentences. Activation profiles of Syntax and Semantics voxels to the 10 mini-sentences conditions from Task 2. Frontal Glasser language regions are presented.

